# Greenhouse conditions in lower Eocene coastal wetlands? – Lessons from Schöningen, Northern Germany

**DOI:** 10.1101/2020.04.24.059345

**Authors:** Olaf K. Lenz, Walter Riegel, Volker Wilde

**Affiliations:** General Directorate, Senckenberg Society for Nature Research, Frankfurt am Main, Germany; Department Palaeontology and Historical Geology, Senckenberg Research Institute and Natural History Museum, Frankfurt am Main, Germany

## Abstract

The Paleogene succession of the Helmstedt Lignite Mining District in Northern Germany includes coastal peat mire records from the latest Paleocene to the middle Eocene at the southern edge of the Proto-North Sea. Therefore, it covers the different long- and short-term climate perturbations of the Paleogene greenhouse. 56 samples from three individual sections of a lower Eocene seam in the record capture the typical succession of the vegetation in a coastal wetland during a period that was not affected by climate perturbation. This allows facies-dependent vegetational changes to be distinguished from those that were climate induced. Cluster analyses and NMDS of well-preserved palynomorph assemblages reveal four successional stages in the vegetation during peat accumulation: (1) a coastal vegetation, (2) an initial mire, (3) a transitional mire, and (4) a terminal mire. Biodiversity measures show that plant diversity decreased significantly in the successive stages. The highly diverse vegetation at the coast and in the adjacent initial mire was replaced by low diversity communities adapted to wet acidic environments and nutrient deficiency. The palynomorph assemblages are dominated by elements such as *Alnus* (Betulaceae) or *Sphagnum* (Sphagnaceae). Typical tropical elements which are characteristic for the middle Eocene part of the succession are missing. This indicates that a more warm-temperate climate prevailed in northwestern Germany during the early lower Eocene.

## Introduction

The long-term warming trend of the early Paleogene greenhouse climate culminated in the Early Eocene Climatic Optimum (EECO) between *c.* 52 and 50 Ma before present (BP) [1]. It was interrupted by short-term warming events, the most prominent being the Paleocene-Eocene Thermal Maximum (PETM or ETM-1, e.g., [2–4]), which is associated with rapid temperature increases at the transition between the Paleocene and Eocene, which is estimated to have lasted about 170 (± 30) kyr [5–7]. Other short-term events followed, such as the ETM-2 (*c*. 53.6 Ma BP [8, 9]) and the ETM-3 (X- or K-event; *c*. 52.5 Ma BP [10, 11]), but did not reach the temperatures of the PETM.

Recognition of these thermal events is has originally been based on carbon isotope excursions (CIE) observed in deep sea cores (e.g., [2,8,10,12–15]) and considered to result from massive release of light carbon (C13) to the atmosphere (e.g., [16, 17]). The effect of the subsequent temperature increase on terrestrial environments has been debated and discussed with respect to migration of biota and extinction of species mainly in the North American Continental Interior Especially the PETM was associated with large scale migrations of mammals including new modern type taxa along open routes in the North American Continental Interior [18, 19]. But reports on corresponding effects of changes in vegetation are ambiguous, if not controversial, and appear to vary considerably depending on regional or local conditions. While species turnovers may have been more prevalent only in the tropics [20], a change from a mesophytic to a more thermophilic flora has been noted in the Bighorn Basin, Wyoming, USA [21]. In mid-latitude locations of Europe vegetation changes occurred essentially on a quantitative basis depending on local conditions or in response to increased precipitation [22, 23]. In order to develop a more coherent picture of the effects of the early Eocene thermal events on the terrestrial vegetation, additional case studies may be necessary.

The sedimentary succession of the former Helmstedt Lignite Mining District, which includes the mines at Schöningen, covers the entire Paleogene greenhouse phase and its gradual demise from the latest Paleocene to the middle Eocene almost continuously in the Helmstedt Embayment at the southern edge of the Proto-North Sea [24–26]. This offers the unique opportunity to trace the effects of all the long- and short-term climate perturbations on Paleogene terrestrial ecosystems across more than 10 million years. The study is part of a current project on changes in composition of the vegetation and plant diversity in the coastal environment of the Helmstedt Embayment across the EECO and its short-term perturbations such as the PETM and ETM-2 by using pollen and spores as proxies.

A first set of isotope analyses from the lower part of the lower Eocene Schöningen Formation revealed a carbon isotope excursion (CIE) from the very top of Seam 1 to the middle of Seam 2 including Interbed 2, indicating a warming event for this interval [27]. However, it is important to note that Seam 1 has been deposited during a period without any perturbations in climate and changes in vegetation were controlled by other factors than climate such as natural succession due to peat aggradation. Therefore, we selected Seam 1, for which three individual sections were available from the Schöningen outcrops and studied them palynologically including multivariate statistical analyses and biodiversity measures to determine more precisely the composition and variability of the regional flora unaffected by warming events. In this way we expect to be able to better identify possible thermophilic elements which may have invaded the area during warming events and to assess concomitant changes in diversity and quantitative composition.

The lignites of the Helmstedt Lignite Mining District, including Seam 1 at Schöningen, are somewhat unique among Eocene lignites in their close control by sea level dynamics and floristically different from the better known and widespread younger Miocene lignites. For images of the field situation at Schöningen we refer to [25, 28].

## Geological setting

The Helmstedt Lignite Mining District is situated within the Paleogene Helmstedt Embayment, which represented the mouth of an estuary opening towards the Proto-North Sea (Fig 1) between major uplifts corresponding to the Harz Mountains to the South and the Flechtingen Rise to the North [29]. The estuary extended far inland towards the area of Halle and Leipzig (Leipzig Embayment [30, 31]). Due to the interaction between changes in sea level, salt withdrawal in the subsurface and climate-related changes in runoff from the hinterland, the area of Helmstedt and Schöningen was subject to frequent changes between marine and terrestrial conditions, repeatedly leading to peat formation [25, 32].

**Fig 1.**
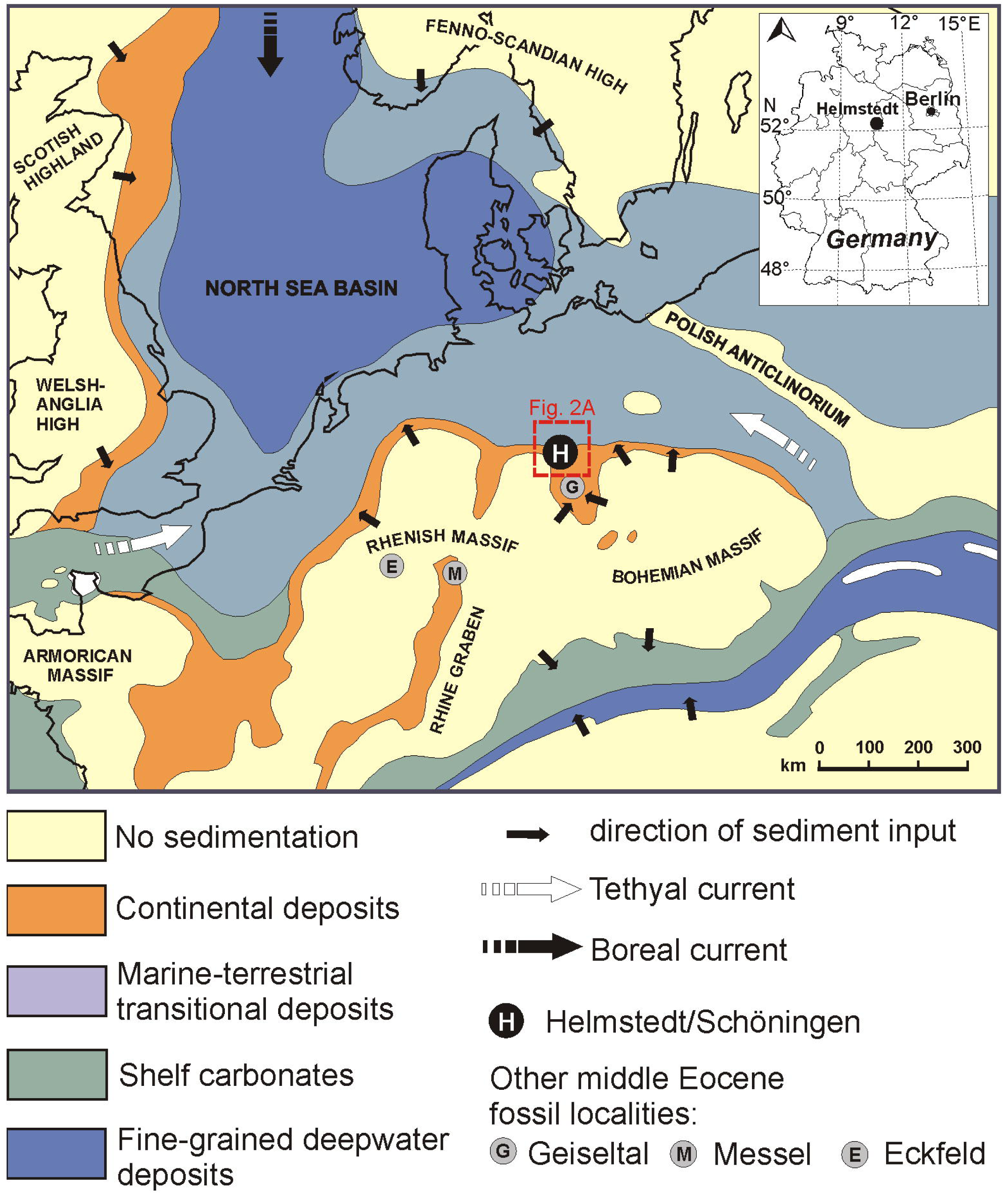
Paleogeographic map of Northwestern Europe during the lower Eocene. The map shows the Helmstedt Embayment at the southern edge of the Proto-North Sea (H) in relation to important middle Eocene fossil localities in Germany, such as the Geiseltal (G), Messel (M), and Eckfeld (E); adapted from [27, 33].

Today, the Paleogene deposits of the Helmstedt Lignite Mining District are limited to two marginal synclines accompanying the more than 70 km long salt wall of Helmstedt-Staßfurt (Fig 2) [29, 34]. Both of the synclines are strongly asymmetric with steeply inclined strata dipping away from a narrow core of Zechstein salt while they are gently dipping on the opposite flanks. The influence of salt-withdrawal on sediment accumulation is indicated by the fact that the maximum thickness of the two lignite bearing sequences and of the individual coal seams moved towards the salt-wall with time [29, 34].

**Fig 2.**
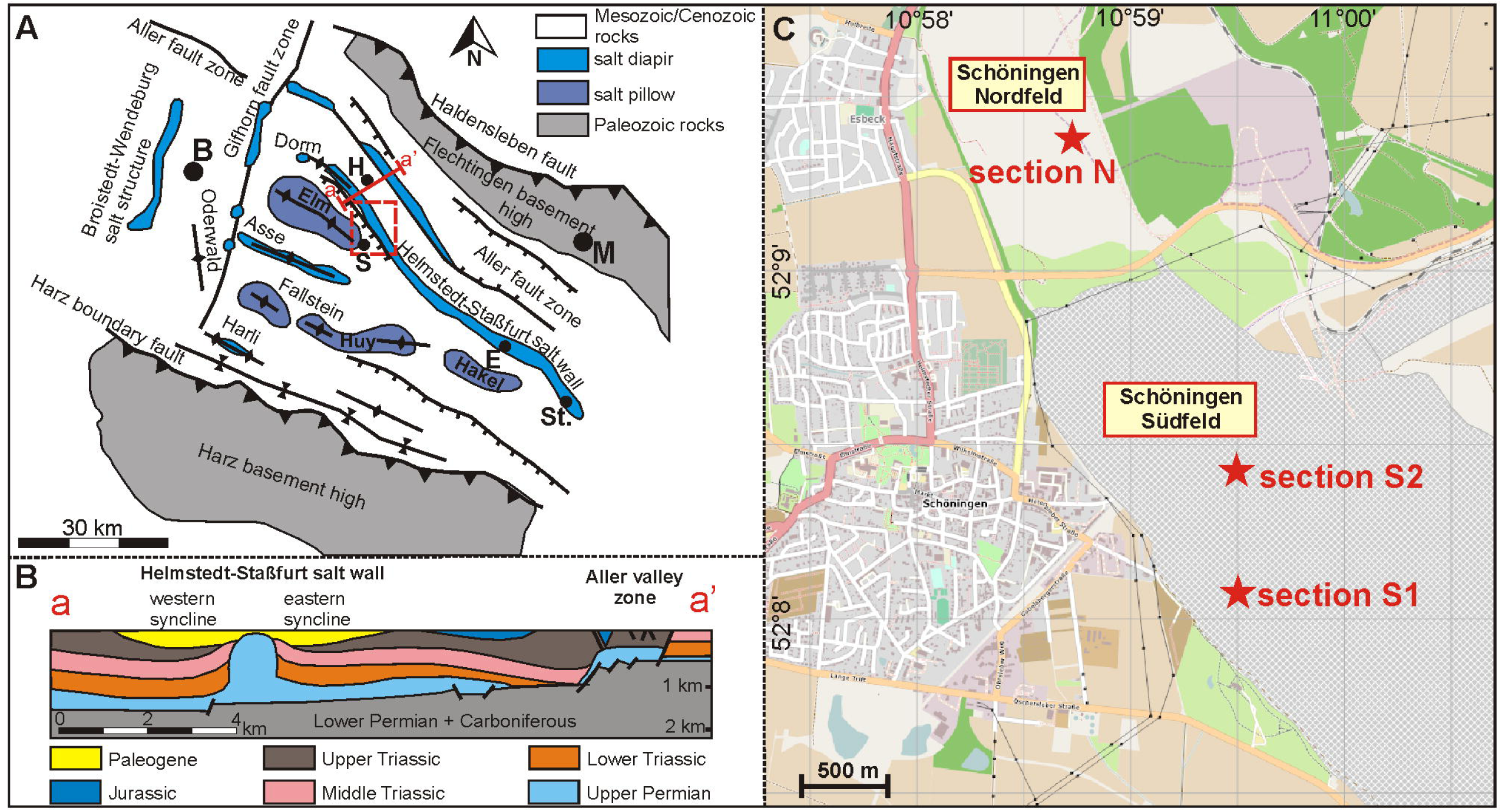
The regional setting of the Helmstedt Lignite Mining District. (A) Geologic structural map of the area between the major uplifts of the actual Harz Mountains to the South and the Flechtingen Rise in the north (modified after [29]). The salt pillows and diapirs are not exposed at the surface, but buried under Mesozoic and Cenozoic sedimentary rocks. The red frame marks the detail presented in (C) (B = Braunschweig, H = Helmstedt, S = Schöningen, E = Egeln, St = Staßfurt). (B) Cross-section through the study area, showing the Helmstedt-Staßfurt salt wall and related synclines (modified after [29]). (C) The former opencast mines Schöningen Nordfeld and Schöningen Südfeld east of Schöningen. The positions of the three studied sections of Seam 1 are indicated. The map has been rendered with the software Maperitive using geodata from OpenStreetMap.

An approximately 400 m thick Paleogene succession in both synclines unconformably rests upon Mesozoic sediments of Triassic and lower Jurassic age (Figs 2 and 3) [29, 35]. The position of the lignites and two major marine transgressions in the sequence suggested a subdivision of the Paleogene strata from bottom to top in underlying sediments (now Waseberg Formation), Lower Seam Group (now Schöningen Formation), Emmerstedt Greensand (now Emmerstedt Formation), Upper Seam Group (now Helmstedt Formation) and overlying marine strata (now Annenberg-, Gehlberg-, und Silberberg Formations) (Fig 2) [25,36,37].

**Fig 3.**
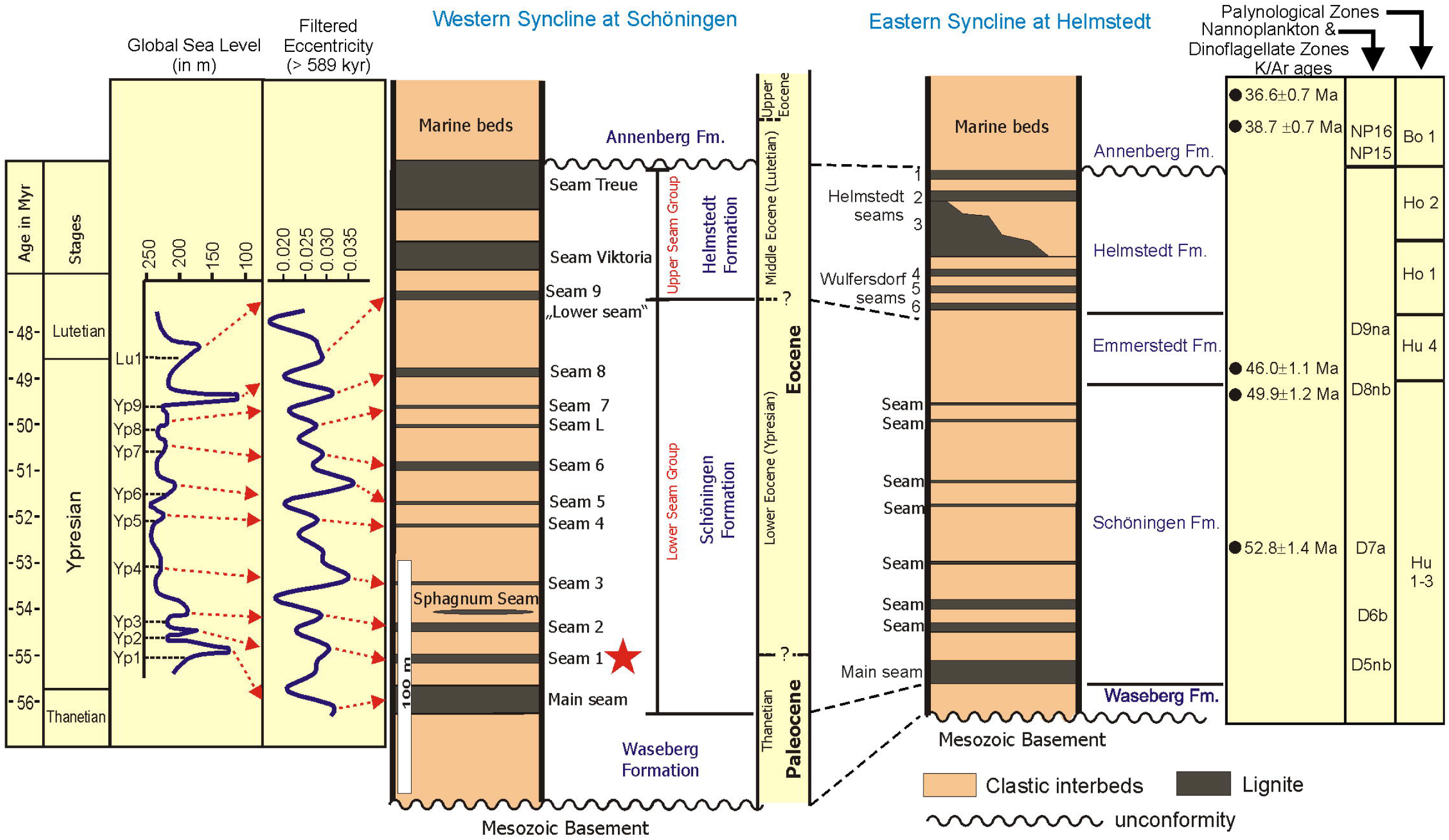
Stratigraphic scheme of the Paleogene succession in the western and eastern Syncline at Helmstedt and Schöningen. The age model for the succession is based on K/Ar-ages [36, 38], nannoplankton zones [36], dinoflagellate zones [39] and palynological zones [40, 41]. Data for global changes in Paleogene sea-level [42] and higher order orbital cyclicity (long eccentricity >589 kyr) [43] are used for a putative correlation to seams in the Schöningen Südfeld section. The asterisk points to the stratigraphic position of the studied sections

The conventional age model (Fig 3) for the coal-bearing part of the Paleogene succession of the Helmstedt Lignite Mining District is mainly based on scattered radiometric ages from glauconites [36, 38] as well as biostratigraphic data from nannoplankton [36], dinocysts [38, 39] and palynomorphs [40, 41]. The data for the Schöningen, Emmerstedt and Helmstedt formations were mostly derived from wells near Helmstedt in the eastern Syncline and have simply been transferred by lithologic correlation to both synclines in the rest of the area. They suggest a lower Eocene (Ypresian) age for the Schöningen Formation and a middle Eocene (Lutetian) age for the Helmstedt Formation.

More recent results on quantitative data for the dinoflagellate cyst genus *Apectodinium* and carbon isotopes from the section at Schöningen in the western syncline indicate that the lowermost part of the Schöningen Formation may still be of Paleocene age [25, 27]. Furthermore, this age model appears consistent when the succession of seams at Schöningen is compared to global changes in Paleogene sea-level and higher order cyclicity [25].

The Schöningen Formation as exposed in the opencast mine Schöningen-Südfeld (Fig 2) of the western syncline has a thickness of about 155 m, including 9 almost continuous seams (Main Seam and Seam 1 (this study) to Seam 9) and some additional seams of limited extent, including Seam “L” and the “*Sphagnum* Seam” [25, 44]. The Emmerstedt Formation cannot be identified at Schöningen since the characteristic greensand is missing.

Due to a lack of radiometric dates and relevant biostratigraphic information, the exact position of both, the Paleocene-Eocene boundary and the Ypresian-Lutetian boundary still remain unknown at Schöningen. The frequency of *Apectodinium* spp. just above Seam 1 in the western syncline [25, 27] has been discussed as indicating the Paleocene-Eocene Thermal Maximum (PETM) (see [45–51]). However, the marker species of the PETM in open marine environments *A. augustum* [45], now *Axiodinium augustum* [52], is not found among the countless *Apectodinium* cysts above Seam 1 [27]. Furthermore, since *Apectodinium* acmes occur in marginal marine areas in the North Sea basin also at other times during the early and middle Eocene [53] the distinct carbon isotope excursion (CIE) co-occurring with the *Apectodinium* acme in Interbed 2 cannot unequivocally be related to the PETM and may point to another later warming event [27]. Thus, Seam 1, which is in the focus of our study, cannot be unambiguously dated at the moment.

## Methods

### Sampling and sample processing

Three sections of the 2.75 m to 3.75 m thick Seam 1 have been studied (Figs 2C and 4) in now abandoned opencast mines in the western syncline at Schöningen. Access to the sections in the field was provided by the Helmstedter Revier GmbH (formerly Braunschweigische Kohlen-Bergwerke, BKB and later E.ON). Section N (14 samples) was located in mine Schöningen-Nordfeld (52°09’23.8"N 10°58’44.0"E), while the sections S1 (12 samples) and S2 (30 samples) were taken in mine Schöningen-Südfeld (S1: 52°08’07.1"N 10°59’29.7"E; S2: 52°08’27.9"N 10°59’24.5"E). The field situation for S2 has been documented in [28]. The palynological data of sections S1 und S2 are based on new quantitative counts while the analysis of section N is based on data of Hammer-Schiemann (1998, unpublished doctoral dissertation, University of Göttingen).

**Fig 4.**
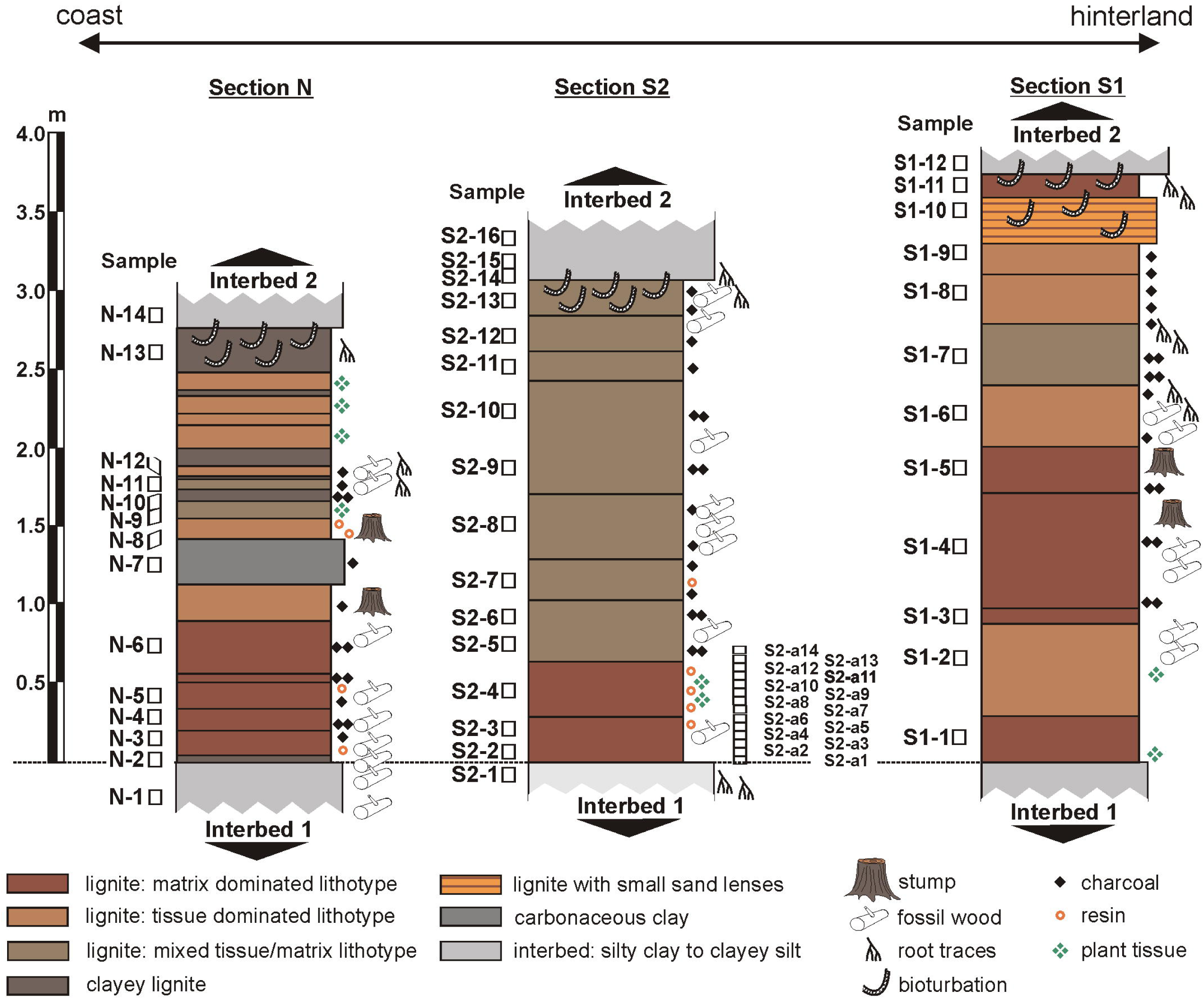
Lithological logs of the three studied sections of Seam 1. Section N is located in mine Schöningen Nordfeld, sections S1 and S2 are from mine Schöningen Südfeld. Information on clastic sediment and lignite texture is based on field observation. Numbers indicate palynological samples.

Most of the individual layers which have been distinguished within Seam 1 in the three sections are represented in the present study by a single sample at least. In order to include the interbed/seam transitions, seven samples from the underlying Interbed 1 and the succeeding Interbed 2, which are both mainly composed of silty clays to clayey silts, have been studied in addition (Fig 4).

Palynological preparation followed the standard procedures as described by [54] including the successive treatment with hydrofluoric acid (HF), hydrogen peroxide (H_2_O_2_) and potassium hydroxide (KOH). Flocculating organic matter was removed by briefly oxidizing the residue with hydrogen peroxide (H_2_O_2_) after sieving with a mesh size of 10 µm. Remaining sample material and slides are stored at the Senckenberg Forschungsinstitut und Naturmuseum, Sektion Paläobotanik, Frankfurt am Main, Germany (repository number Schö XXXIII). No permits were required for the described study, which complied with all relevant regulations.

### Quantitative palynological analysis

Numerical analyses of palynological data are based on quantitative palynomorph counts. At least 300 individual palynomorphs per sample were identified and counted at 400 times magnification to obtain a representative dataset for statistical analysis. A complete list of all palynomorphs encountered during the present study with full names including authors is presented in the taxonomic list (S1 Table). Furthermore, raw data values for section N (S2 Table), section S1 (S3 Table) and section S2 (S4 Table) are presented in the appendix. Identification of palynomorphs is based on the systematic-taxonomic studies of [24,55–58].

Despite the good preservation of palynomorphs 5-10% of the total assemblages could not be identified and have been counted as "Varia". To minimize potential errors in identification and counting of individual species some morphologically similar taxa have been lumped in the pollen diagrams and for statistical analysis, such as different species of the genera *Triporopollenites* or *Triatriopollenites*. In total, 45 groups of palynomorphs have been distinguished (see S1 Appendix). To obtain a robust data set for diversity analyses the slides from 11 lignite samples of section S1 were additionally scanned for rare taxa that were not recorded during routine counting.

The pollen diagrams show the abundance of the most important palynomorphs as percentages. They are arranged according to their weighted average value (WA regression) [59] in relation to depth by using the software C2 1.7.6 [60]. However, WA regression leads to a different arrangement of taxa in the three pollen diagrams, since they may have different ranks in each of the three data sets. Therefore, we used the median rank of the individual taxa to get an identical arrangement of palynomorphs in the three pollen diagrams. Pollen and spores were calculated to 100%, whereas algae, such as *Botryococcus*, dinoflagellates and other organic particles, such as fungal remains, cuticles or charcoal were added as additional percentages (in % of the total sum of pollen and spores).

### Statistical analysis

Statistical analyses followed a routine which has already been applied by the authors in previous studies [61, 62]. We used Wisconsin double standardized raw data values [63–66]. Wisconsin standardization scales the abundance of each taxon or group of taxa to their maximum values and represents the abundance of each of these palynological variables by its proportion in the sample [67]. This equalizes the effects of rare and abundant taxa and removes the influence of sample size on the analysis [63, 64].

For the robust zonation of the pollen diagrams of the three sections and to identify samples with similar palynomorph contents, Q-mode cluster analysis was established using the unweighted pair-group average (UPGMA) method and the Bray-Curtis distance (software PAST 3.26 [68]). Furthermore, to illustrate compositional differences and ecological trends in Seam 1, and to visualize the level of similarity between samples, non-metric multidimensional scaling (NMDS) with the standardized raw data values and the Bray-Curtis dissimilarity [63, 69] has been performed for each of the three studied sections as well as for the complete data set using the software PAST 3.26 [68]. NMDS is the most robust unconstrained ordination method in ecology [70] and has been successfully applied to palynological data in previous studies (e.g., [61,67,71-73]. It avoids the assumption of a linear or unimodal response model between the palynomorph taxa and the underlying environmental gradients as well as the requirement of normal distributed data.

### Diversity analysis

In addition to the quantitative analysis of the 45 groups of palynomorphs that are presented in the pollen diagrams, in section S1 the palynomorph assemblage has been studied with the highest possible taxonomic resolution allowing a detailed analysis of the diversity of the microflora (S5 Table). For diversity analysis, morphologically distinct pollen “species” were recorded representing the morpho-diversity of the palynomorph assemblage. However, these morpho-types do not necessarily reflect different parent plants and may also include morphological variation within the same plant family or genus. Furthermore, morphological diversity within a natural species may include different morpho-types [74]. Nevertheless, since this affects all samples to the same extent, the diversity measures still lead to a robust picture of the diversity of the parent vegetation.

To estimate the changes in taxonomic diversity between single samples and different pollen zones (PZs) within Seam 1, several calculations for species richness and evenness were applied, using tools for biodiversity analysis as provided by [75, 76]. Richness is simply the number of taxa within an ecosystem, which is here calculated as the total number of palynological taxa within a sample or a PZ [77]. It can be measured at different scales, for which mainly the three terms alpha, beta, and gamma diversity have been used [78]. The definitions of these terms were originally based on the comparison of diversity in different areas or regions. Here, we use these terms to describe the temporal comparison of diversity changes within Seam 1. Furthermore, we use the term point diversity (within-sample diversity resp. standing richness of [79]) for the richness within a single sample which reflects the total number of taxa as found in the counted number of individual grains [80]. Alpha diversity is regularly related to the diversity within a community or habitat [80] and is here used as a measure for diversity within a PZ, since this represents a specific community during the evolution of the vegetation in Seam 1. Gamma diversity normally includes the species richness in a larger area within a landscape [80] but is here used as a measure for the richness in the complete seam summarizing the vegetation of the peat-forming communities at Schöningen. Beta diversity that links alpha and gamma diversities is here used as a measure of the difference in species composition between two samples, within a specific PZ or within the whole seam [78,80–82]. Here we adapt Whittaker’s [78, 83] original suggestion for calculating beta diversity, which is most frequently employed in ecological studies [84]. For comparison between two samples, beta diversity is calculated by the total number of species within the two samples divided by the average species number within the two samples. Beta diversity calculations within a PZ and within Seam 1 are calculated as the total species number within the specific PZ or the whole seam divided by the average species number in samples from the PZ/Seam 1. We applied software PAST 3.26 [68] for calculation of point and beta diversity as well as EstimatesS v. 9.1.0 [76] for the analysis of alpha and gamma diversity.

Species richness cannot directly be estimated by observation and not accurately measured, because the observed number of species in a sample is always a downward-biased estimator for the complete species richness in an assemblage [85]. Therefore, the calculation of the number of palynological species within a single sample or a PZ in the succession of Seam 1 is always an underestimate of the possible number of species. Nevertheless, the calculated richness values can be used as reliable information at least on relative changes of point and alpha diversity.

Evenness is the distribution of pollen taxa within a pollen assemblage [80]. A low evenness indicates an assemblage with one or more dominant taxa, characterized by high numbers of pollen grains of the same types, whereas high evenness points to an assemblage without dominant taxa, indicated by equally distributed taxa [86]. Evenness (E) has been calculated using the formula provided by [77] (E= H/ln(R)) producing evenness values between 0 (low evenness) and 1 (high evenness). For Shannon-Wiener index (H) and richness (R) we used the estimations provided by [75] based on calculations for point diversity within 300 counts, for alpha diversity within 5 samples and for gamma diversity within 20 samples (Tables 1 and 2).

**Table 1:** Estimations of palynological richness and evenness based on individual rarefaction analysis for 11 lignite samples of section S1 (point diversity).

|  | S1-1 | S1-2 | S1-3 | S1-4 | S1-5 | S1-6 | S1-7 | S1-8 | S1-9 | S1-10 | S1-11 |
| --- | --- | --- | --- | --- | --- | --- | --- | --- | --- | --- | --- |
| Individuals counted | 416 | 346 | 343 | 502 | 305 | 316 | 304 | 316 | 310 | 329 | 328 |
| S<sub>obs</sub> | 74 | 88 | 63 | 59 | 30 | 41 | 48 | 45 | 50 | 51 | 72 |
| S(est)<sub>300</sub> | 61.7 | 81.6 | 58.6 | 46.3 | 29.7 | 40.0 | 47.8 | 43.8 | 49.3 | 48.3 | 68.2 |
| S(est)<sub>300</sub> Std.err, 2σ, Lower | 56.1 | 77.3 | 54.9 | 40.8 | 28.7 | 38.1 | 46.8 | 41.7 | 47.7 | 45.3 | 64.7 |
| S(est)<sub>300</sub> Std.err, 2σ, Upper | 67.4 | 86.0 | 62.3 | 51.7 | 30.8 | 41.9 | 48.7 | 45.9 | 50.9 | 51.3 | 71.7 |
| Shannon-Wiener index | 3.24 | 3.70 | 3.13 | 2.84 | 2.19 | 2.76 | 3.13 | 2.72 | 3.02 | 2.37 | 3.19 |
| Evenness (E) | 0.79 | 0.84 | 0.77 | 0.74 | 0.65 | 0.75 | 0.81 | 0.72 | 0.78 | 0.61 | 0.76 |

S<sub>obs</sub>: Actual number of taxa within the samples; S(est)<sub>300</sub>: Expected number of species for 300 counted palynomorphs using the algorithm of [92]; S(est)<sub>300</sub> Std.err, 2σ: Lower and upper bounds for a standard error of two-sigma of unconditional variance for 300 palynomorphs; evenness calculation using the method of [77]: $E = \text{Shannon-Wiener index} / \ln(S_{\text{obs}})$ .

## Results

The three studied sections in the western syncline have been arranged from NW to SE following the order of proximity to the sea (Fig 2C). Section N is closest to the sea and the only one including some horizons of clayey lignite and carbonaceous clay (Fig 4). Four lithotypes of lignite have been distinguished in the field, depending mainly on the relative proportion of fine-grained matrix and tissue remains as well as clay content, i.e., matrix dominated, tissue dominated, mixed tissue/matrix lithotype and clayey lignite (Fig 4). The terminology is adapted from [87]. Matrix dominated lithotypes tend to occur preferentially in the lower part of the three sections. However, despite a general upward increase, the degree of tissue preservation remains relatively low throughout most of the seam. In many sections the top of Seam 1 is intensely bioturbated. Charcoal shows a general upward increase in the more inland sections S1 and S2, but no correlation with specific lithotypes.

Based on unconstrained Q-mode cluster analysis (Figs 5B, 6B, 7B) five distinct palynomorph assemblages have been recognized in the three studied sections, which can be distinguished by NMDS (Figs 5C, 6C, 7C). They are arranged as palynozones (PZ) in a vertical succession, which shows that the general development of the vegetation was identical in the three sections of Seam 1. The term “palynozone” is used here in the sense of representing plant communities [88]. Abundance variations of palynomorphs within PZs between sections indicate local differences in vegetation patterns.

**Fig 5.**
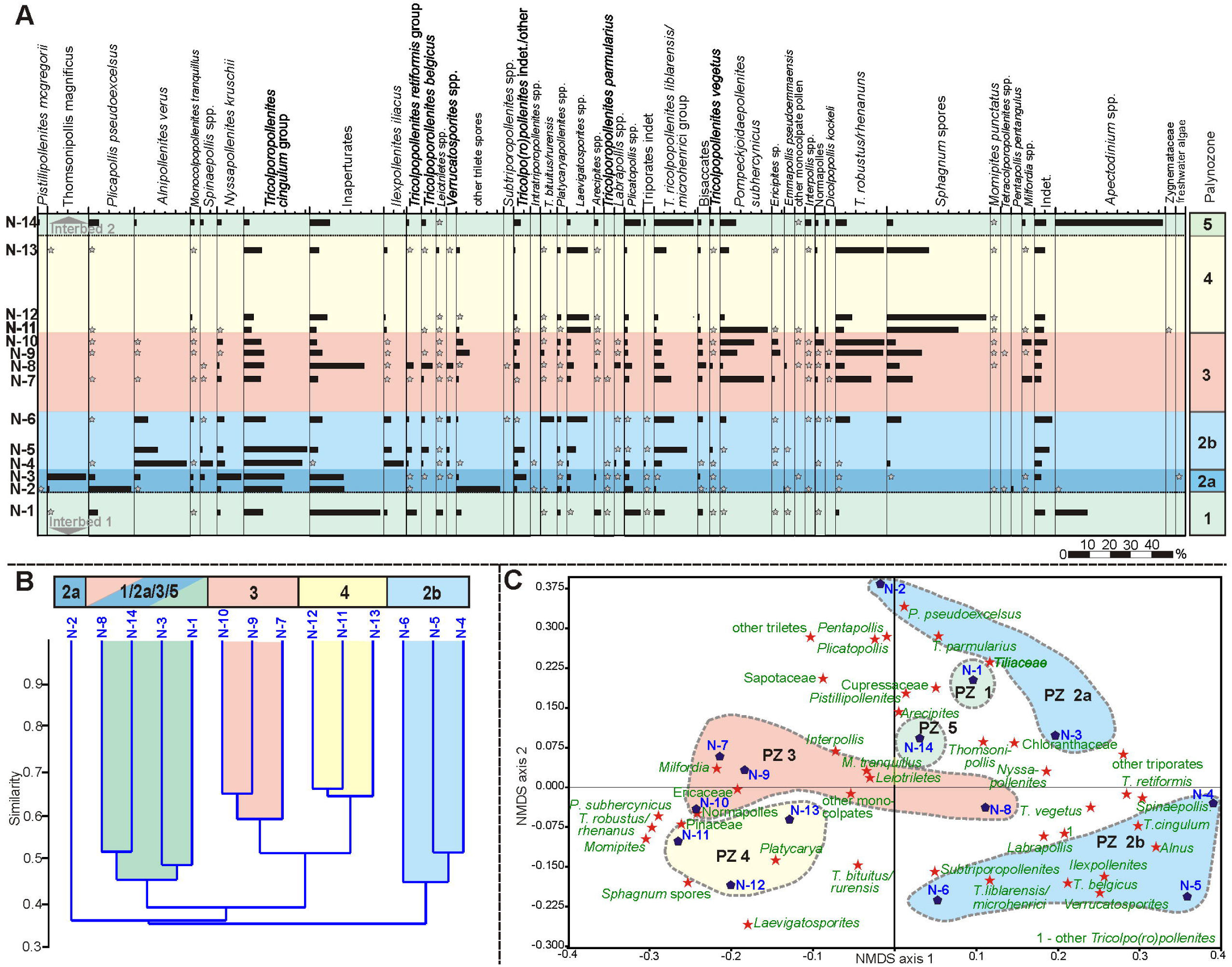
Pollen diagram, cluster analysis and NMDS of section N. (A) Pollen diagram of 14 samples from the top of Interbed 1 to the base of Interbed 2 of section N showing the most common palynomorph taxa. The zonation in different PZs is based on cluster analysis (B) Result of an unconstrained cluster analysis of Wisconsin double standardized raw-data values using the unweighted pair-group average (UPGMA) method together with an Euclidean distance (C) Non-metric multidimensional scaling (NMDS) plot of palynological data using the Bray-Curtis dissimilarity and Wisconsin double standardized raw-data values. The scatter plot shows the arrangement of samples and palynomorph taxa.

**Fig 6.**
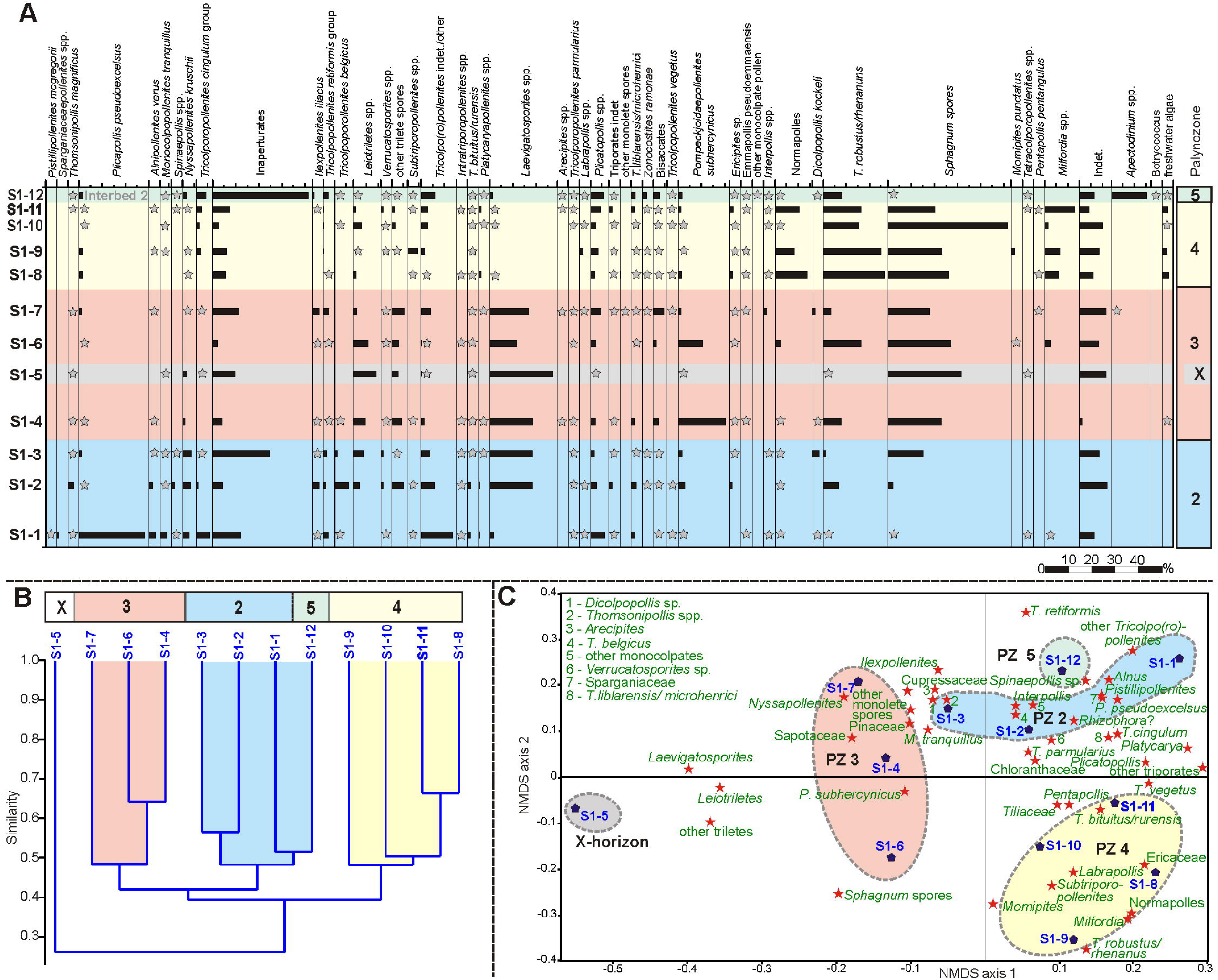
Pollen diagram, cluster analysis and NMDS of section S1. (A) Pollen diagram of 12 samples from the base of Seam 1 to the base of Interbed 2 of section S1 showing the most common palynomorph taxa. The zonation in different PZs is based on cluster analysis (B) Result of an unconstrained cluster analysis of Wisconsin double standardized raw-data values using the unweighted pair-group average (UPGMA) method together with an Euclidean distance (C) Non-metric multidimensional scaling (NMDS) plot of palynological data using the Bray-Curtis dissimilarity and Wisconsin double standardized raw-data values. The scatter plot shows the arrangement of samples and palynomorph taxa.

**Fig 7.**
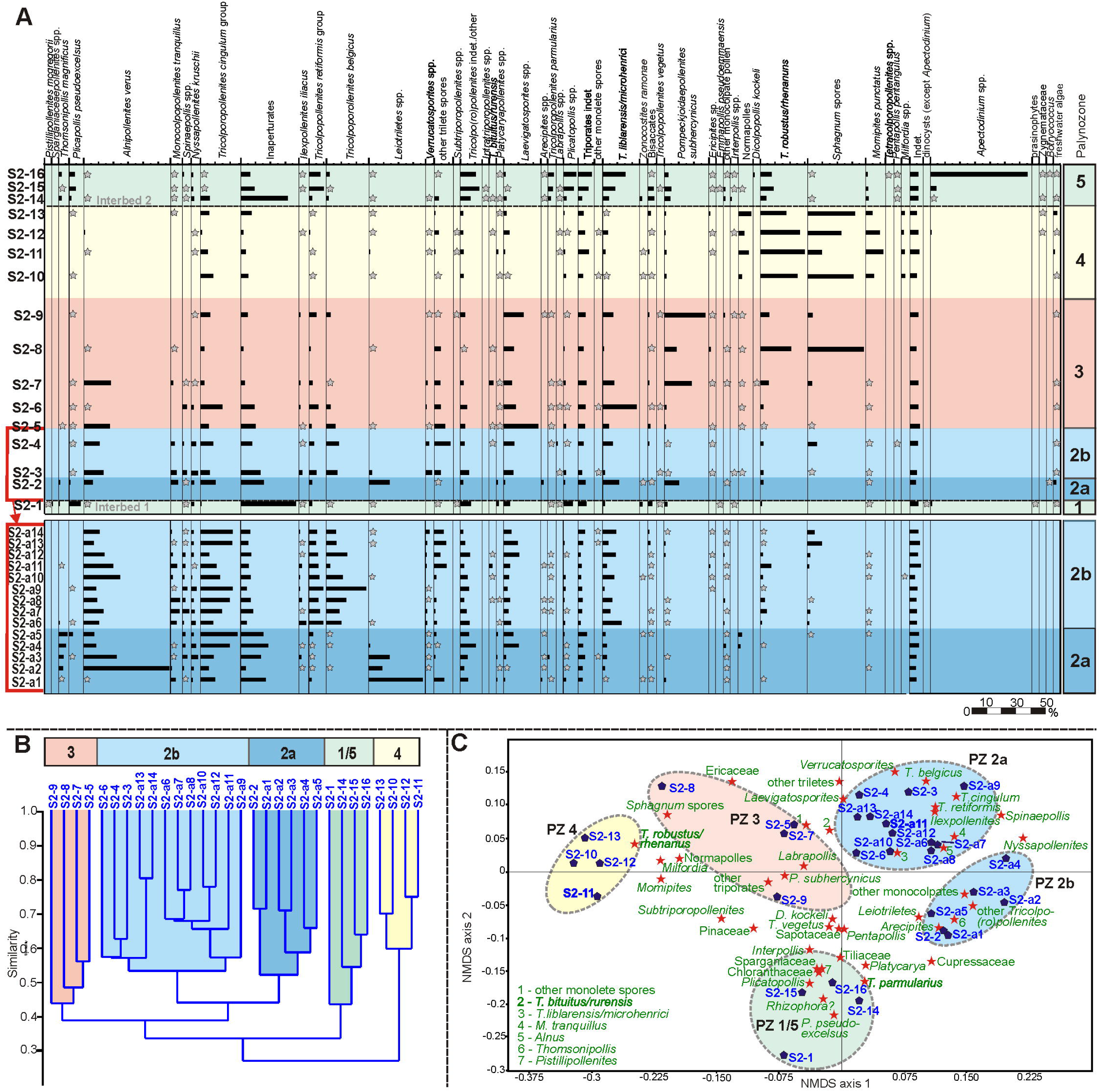
Pollen diagram, cluster analysis and NMDS of section S2. (A) Pollen diagram of 30 samples from the top of Interbed 1 to the base of Interbed 2 of section S2 showing the most common palynomorph taxa. The zonation in different PZs is based on cluster analysis (B) Result of an unconstrained cluster analysis of Wisconsin double standardized raw-data values using the unweighted pair-group average (UPGMA) method together with an Euclidean distance (C) Non-metric multidimensional scaling (NMDS) plot of palynological data using the Bray-Curtis dissimilarity and Wisconsin double standardized raw-data values. The scatter plot shows the arrangement of samples and palynomorph taxa.

PZ 1 and PZ 5 include samples from the adjacent Interbeds 1 and 2, and reflect the state of vegetation during the marine-terrestrial transition below and the terrestrial-marine transition above the seam. PZs 2, 3, and 4, on the other hand, represent different stages of the peat forming vegetation during seam formation.

### Palynozone 1 (top Interbed 1)

The two samples from Interbed 1 (sample N-1, Fig 5A and sample S2-1, Fig 7A) show marine influence with the occurrence of dinocysts (*Apectodinium* spp.). The NMDS of S2 samples (Fig 7C) shows that sample S2-1 is clearly different from the seam, because it is plotted in the ordination space on the negative side of NMDS axis 2 together with samples of Interbed 2 (PZ 5) but separate from all of the lignite samples (PZ 2 - 4). Sample N-1 (Fig 5C) is plotted in the upper right corner of the ordination space very close to samples from the base of Seam 1 (PZ 2a) indicating a more gradual change of the vegetation from marginal marine habitats to the peat-forming environment at this site.

The only true mangrove element *Rhizophora* (*Zonocostites ramonae,* Fig 8I), in the Schöningen Formation, occurs in low abundance in PZ 1 of section S2 and in PZ 5 of sections S1 and S2. *Inaperturopollenites* spp. (Cupressaceae s.l., Figs 8A, B) dominate the pollen assemblages with values of up to 37%. Other common taxa of sample N-1 are *Tricolporopollenites cingulum* (Fagaceae, 9.4%, Figs 8C, D, E), *Plicatopollis* spp. (Juglandaceae, 7.7%, Fig 8K), *Tricolpopollenites liblarensis* (Fagaceae, Fig 8F) and *T. retiformis* (Salicaceae, Fig, 8H), each with 5.1% as well as *Plicapollis pseudoexcelsus* (Juglandaceae?, 4.3%, Fig 8J).

**Fig 8.**
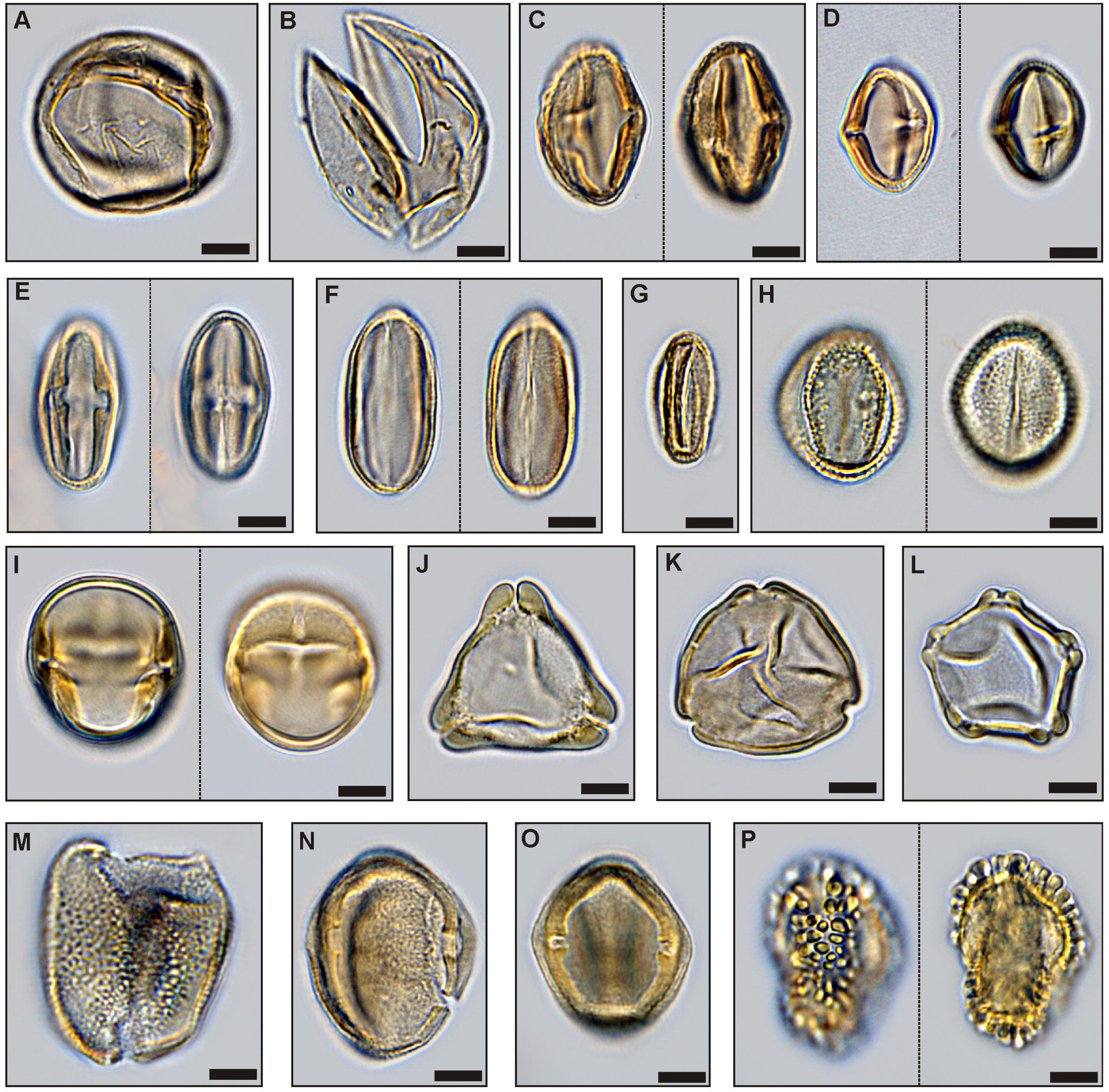
**Important palynomorphs of PZs 1, 2 and 5**. (A) *Inaperturopollenites concedipites,* Cupressaceae s.l. (sample S1-12), (B) *Cupressacites bockwitzensis,* Cupressaceae s.l. (sample S1-12); (C) *Tricolporopollenites cingulum fusus,* Fagaceae (morphotype 1 with a rough exine, larger than morphotype 2; sample S1-12), (D) *Tricolporopollenites cingulum fusus,* Fagaceae (morphotype 2 with a smooth exine, smaller than morphotype 1; sample S1-12), (E) *Tricolporopollenites cingulum pusillus,* Fagaceae (morphotype 2, sample S1-9), (F) *Tricolpopollenites liblarensis liblarensis,* Fagaceae (sample S1-12), (G) *Tricolpopollenites quisqualis,* Fagaceae (sample S1-12); (H) *Tricolpopollenites retiformis,* Salicaceae (sample S1-4); (I) *Zonocostites ramonae,* ?Rhizophoraceae (sample S1-8); (J) *Plicapollis pseudoexcelsus,* ?Juglandaceae (sample S1-9); (K) *Plicatopollis hungaricus,* Juglandaceae (sample S1-3); (L) *Alnipollenites verus,* Betulaceae (sample S1-3); (M) *Dicolpopollis kockeli,* Arecaceae (sample S1-3); (N), (O) *Nyssapollenites kruschii accessorius,* Nyssaceae (samples S1-12, S1-3); (P) *Ilexpollenites iliacus,* Aquifoliaceae (sampleS1-4); scale bars: 10µm

In sample S2-1 *Plicatopollis* spp. (6.7%) and *P. pseudoexcelsus* (8.0%) are also very common, while the fagaceous taxa *T. liblarensis* and *T. cingulum* as well as *T. retiformis* are less frequent compared to sample N-1.

### Palynozone 2 (Seam 1)

PZ 2 (Figs 5A, 6A, 7A) includes the lower part of Seam 1 and can be subdivided into two subzones in sections N and S2. The difference to other samples of the seam is best expressed in section S2 in the NMDS where samples of PZ 2 are clearly separate on the right side of the ordination space (Fig 7C). In the other two sections, samples from adjacent interbeds are close to PZ 2 samples in the ordination space, thus, indicating the proximity of PZ 2 to the pollen assemblages of Interbed 1 and Interbed 2 (Fig 5C, resp. 6C).

In contrast to PZ 1, dinoflagellate cysts are completely missing in PZ 2. In the composition of the pollen assemblage the most striking change is the occurrence of *Alnipollenites verus* (Betulaceae, *Alnus,* Fig 8L), which reaches a maximum of 57.5% in section S2. Although much lower, the maxima of *A. verus*, too, occur in PZ 2: 25.3% in section N resp. 2.9% in section S1.

*Tricolporopollenites cingulum* (Fagaceae) is among the dominant elements in sections N and S2. In section S1 maximum values are distinctly lower, but also reached in PZ 2. Other taxa with maxima in PZ 2 are *Spinaepollis spinosus* (Euphorbiaceae?), *Nyssapollenites* spp. (Nyssaceae, Figs 8N, O) and *Ilexpollenites* spp. (Aquifoliaceae, Fig 8P). The lowest values for these taxa occur again in section S1. *Inaperturopollenites* sp. is still characterized by high values (24.6%), a slight decrease, however, from PZ 1.

A few taxa strongly decrease within PZ 2 and, therefore permit the separation of subzones PZ 2a and PZ 2b for sections N and S2. This is the case, in particular, for *Thomsonipollis magnificus* (unknown botanical affinity, Fig 9F) which drops from 18% in section N to near absence in PZ 2b. Similarly, the fern spores *Leiotriletes* spp. (Schizaeaceae, Figs 9B, C, D) and other trilete spores disappear almost completely in PZ 2b except for a slight increase at the top of PZ 2 in section S1. These spores are replaced in PZ 2b by other fern spores such as *Laevigatosporites* spp. (Polypodiaceae, Fig 9 E), which are rare in PZ 2a.

**Fig 9.**
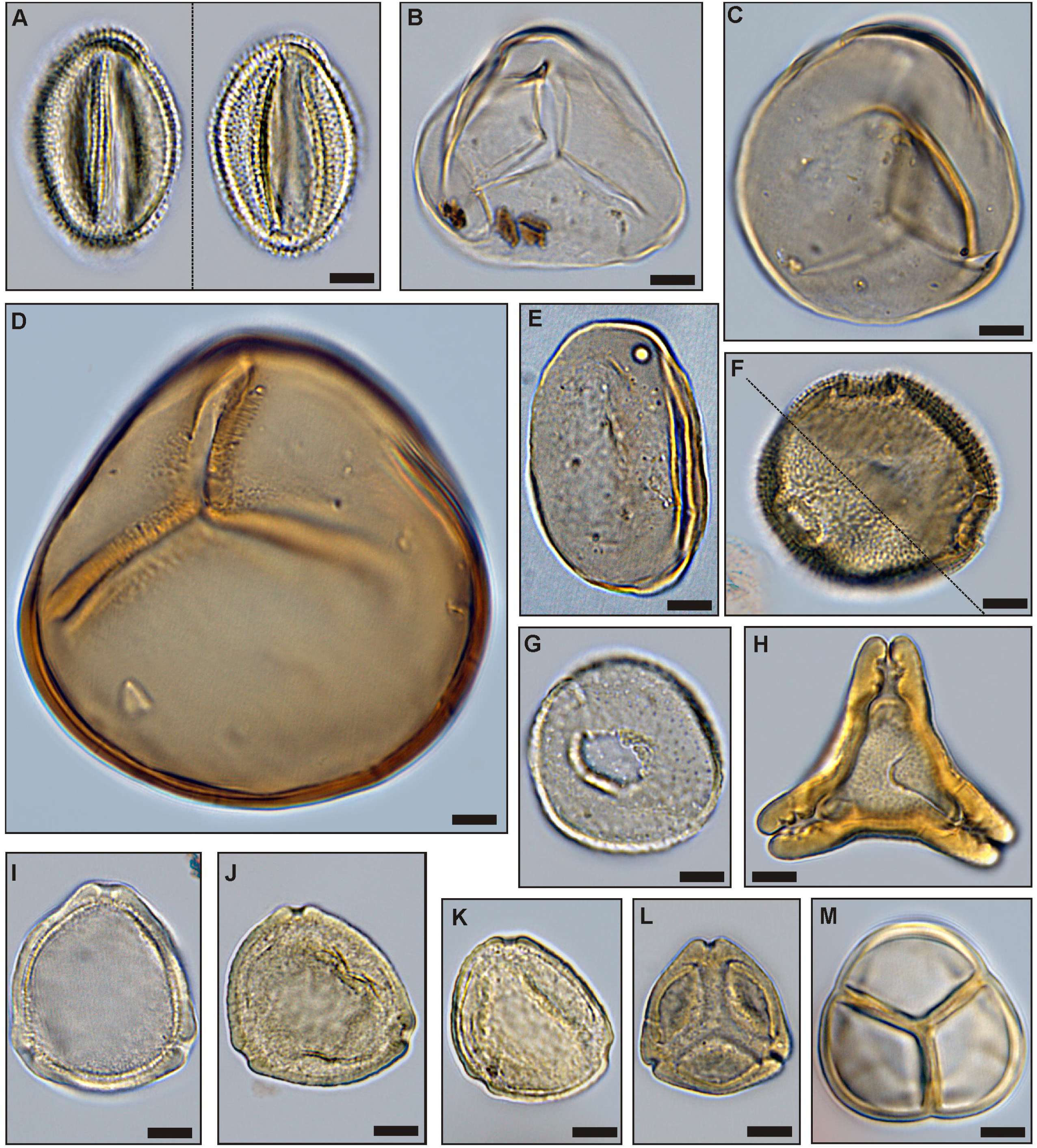
Important palynomorphs of PZs 3 and 4. (A) *Tricolporopollenites belgicus,* unknown botanical affinity (sample S1-2), (B) *Leiotriletes microadriennis,* Schizaeaceae (sample S1-4), (C) *Leiotriletes adriennis,* Schizaeaceae (sample S1-6), (D) *Leiotriletes paramaximus,* Schizaeaceae (sample S1-4); (E) *Laevigatosporites discordatus,* Polypodiaceae (sample S1-3); (F) *Thomsonipollis magnificus,* unknown botanical affinity (sample S1-2); (G) *Milfordia incerta,* Restionaceae (sample S1-9); (H) *Basopollis atumescens,* unknown botanical affinity (sample S1-8); (I) *Triporopollenites crassus,* Myricaceae (sample S1-10), (J) *Triporopollenites robustus,* Myricaceae (sample S1-8), (K) *Triporopollenites rhenanus,* Myricaceae (sample S1-8), (L) *Pompeckjoidaepollenites subhercynicus,* unknown botanical affinity (sample S1-3), (M) *Ericipites ericius,* Ericaceae (sample S1-11); scale bars: 10µm

The relative loss of these taxa in PZ 2b is in part compensated by increases in *Monocolpopollenites tranquillus* (Arecaceae, *Phoenix*), *Tricolpopollenites retiformis, Tricolporopollenites belgicus* (unknown botanical affinity, Fig 9A) and *Tricolpopollenites liblarensis* in section S2 (Fig 7A), in section N additionally by *Ilexpollenites* spp. (Fig 5A). The small number of samples in section S1 here precludes a subdivision of PZ 2.

### Palynozone 3 (Seam 1)

PZ 3 covers the middle of Seam 1 and is represented by 4 samples each in sections N and S1 and by 3 samples in section S2. The NMDS of all sections show that samples of PZ 3 are separated from other PZs in the ordination space indicating a unique assemblage composition. Especially in sections N and S2 (Fig 5C resp. 7C) the samples are plotted midway between those of PZ 2 and PZ 4 indicating that the assemblages include elements from the preceding and succeeding PZs. In section S1 PZ 3 is plotted on the left side of the ordination space clearly separated from the other two seam-related PZs (Fig 6C). However, sample S1-5 is plotted far away from the other samples of PZ 3 on the negative end of NMDS axis 1 (Fig 6C) suggesting a difference in assemblage composition not readily recognized in the pollen diagram (Fig 6A).

*Alnipollenites verus* has virtually disappeared except for local abundance in section S2. Similarly, *Tricolporopollenites cingulum* decreases from relatively high values in section N to 10.8% and from low values in section S1 to less than 1%. Only in section S2 it does remain at similar high levels. Other taxa such as *Spinaepollis spinosus*, *Ilexpollenites* spp., *Nyssapollenites* spp., and *Tricolpopollenites liblarensis* decrease consistently in sections N and S1 as well as *Tricolporopollenites belgicus* in section S2.

*Pompeckjoidaepollenites subhercynicus* (unknown botanical affinity, Fig 9L) suddenly appears with high values (up to 20.2% in section S1) and extends to the base of PZ 4 in sections N and S2. *Triporopollenites robustus* (Myricaceae, Figs 9I - K) as well as spores of Sphagnaceae such as *Sphagnumsporites* sp., *Tripunctisporis* sp. and, *Distancorisporis* sp. [89] (Fig 10) are abundant before becoming prevalent in PZ 4. Both, *T. robustus* and *Sphagnum*-type spores together are already prevalent in PZ 3 of section S1.

**Fig 10.**
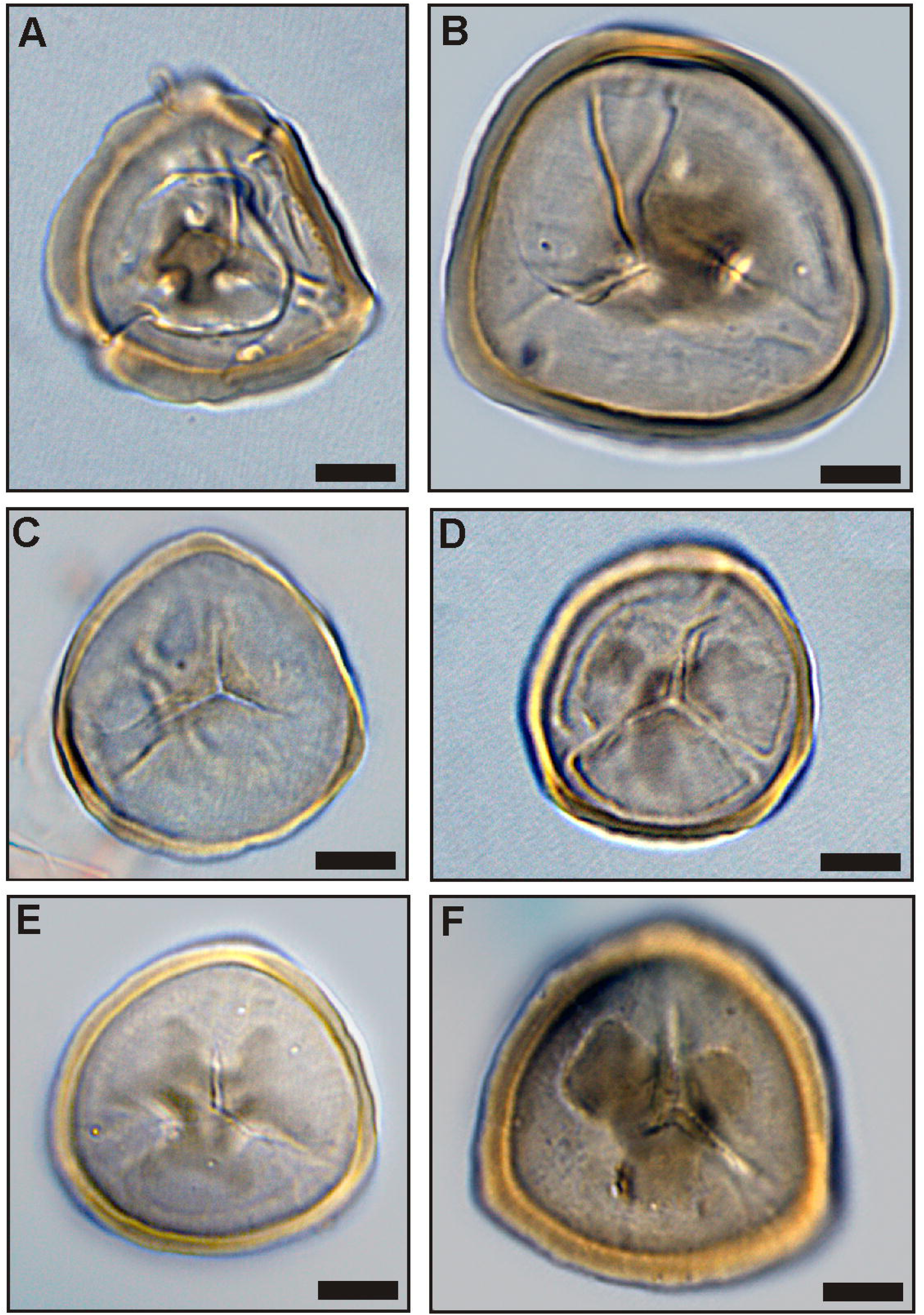
Variation of *Sphagnum*-type spores in PZs 3 and 4. Morphological variation in *Tripunctisporis* sp. (A), (B), *Sphagnumsporites* sp. (C) and *Distancorisporis* sp. (D), (E), (F); scale bars: 10µm

Sample S1-5 (section S1) is special among PZ 3 samples (Fig 6C) due to the high abundance of spores produced by ferns and peat mosses. More than two thirds of the palynomorphs in this sample are composed of spores. Accordingly, *P. subhercynicus* and *T. robustus* remain proportionally rare.

### Palynozone 4 (Seam 1)

PZ 4 comprises the upper part of Seam 1 and represents a significant change from the palynomorph assemblages of preceeding PZs 2 and 3 (Figs 5A, 6A, 7A). This becomes particularly evident in the NMDS. In all three sections samples of PZ 4 are clearly separated from all other samples in the ordination space (Figs 5C, 6C, 7C) due to the dominance of *Sphagnum*-type spores, which reach maximum values between 38% and 52%. A similar dominance is shown for myricaceous pollen, e.g., *Triporopollenites robustus/rhenanus,* with values between 23% and 30%.

*Pompeckjoidaepollenites subhercynicus*, a major element of PZ 3 continues to dominate (up to 28%) into the lower part of PZ 4 in sections N and S2. In section S1, however, it is rare. Pollen of the Normapolles group (e.g. *Basopollis* spp., Fig 9H) and Restionaceae (*Milfordia* spp., Fig 9G) have a strong showing in PZ 4 of section S1, but together with *Ericipites* spp. (Ericaceae, Fig 9M) a distinct reduction over PZ 3 in section N. *Momipites punctatus* (Juglandaceae, *Engelhardia*) is quite common for the first time in section S2 but rare in the other two sections. *Laevigatosporites* spp. are reduced in sections S1 and S2 but increase in section N. *Alnipollenites verus* is extremely rare in PZ 4 of all sections.

### Palynozone 5 (base Interbed 2)

PZ 5 includes mainly samples from Interbed 2 (Figs 5A, 6A, 7A). In all sections a marine influence is indicated by the onset of dinocysts of *Apectodinium* spp. with maximum values of 65.5%. This clearly distinguishes the transition of Interbed 1 to the seam from the transition of the top of the seam to Interbed 2. However, the NMDS of sections N (Fig 5C) and S2 (Fig 7C) show that the palynomorph assemblage composition of both is similar at the beginning and end of seam development. Both PZs are plotted in the ordination space in close vicinity. The NMDS of section S1 (Fig 6C), however, shows similarities in assemblage composition between PZ 5 and PZ 2 since one of the samples (S1-12) is plotted in the NMDS in the upper right corner of the ordination space. But, according to the cluster analysis, the closest similarity of S-12 is to S1-1 at the base of the seam (Fig 6B).

Drastic changes from PZ 4 include the disappearance of *Sphagnum*-type spores and the strong increase of *Inaperturopollenites* spp. with a maximum of 41.3% in section S1. These are similar values as in PZ 1. The pollen of the juglandaceous alliance such as *Plicapollis pseudoexcelsus* (up to 4.8%) and *Plicatopollis* spp. (up to 8.9%) as well as the fagaceous pollen *Tricolpopollenites liblarensis* (up to 15.5%) reach high values that are in the range of their values within PZ 1. *Triporopollenites robustus/rhenanus* (up to 9%) are also common, although the values strongly decrease compared to PZ 4.

### Non pollen/spore palynofacies (section S2)

In section S2 a selection of organic particles, such as fungal remains, periderm cells, cuticle fragments, resin and tannin bodies (resinite/phlobaphinite) as well as charcoal have been quantitatively recorded in addition to palynomorphs and calculated to 100% palynomorphs (Fig 11).

**Fig 11.**
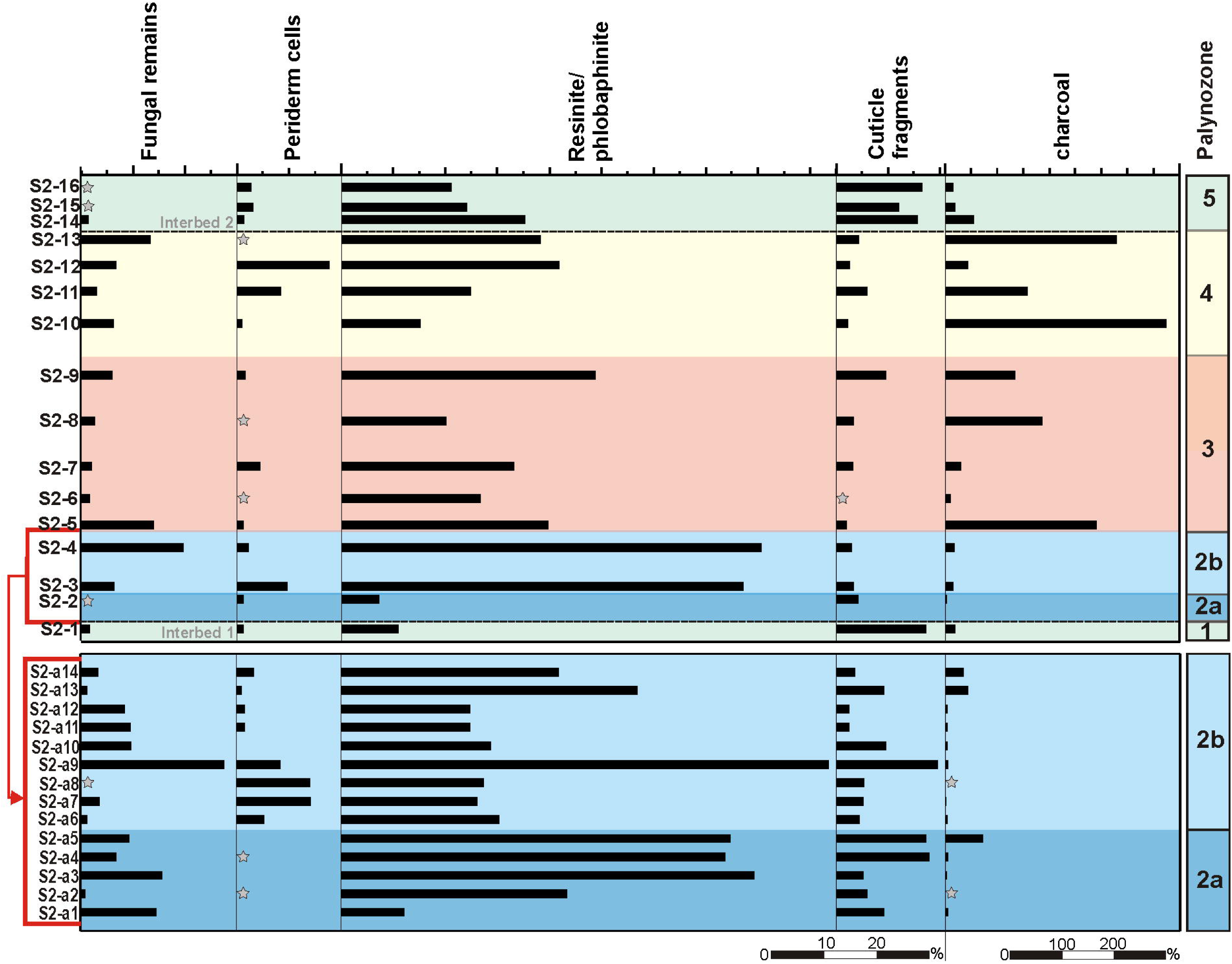
Abundance of non-pollen/spore palynofacies elements (NPP) in section S2. Diagram of 30 samples from the top of Interbed 1 to the base of Interbed 2 of section S2 showing the distribution of NPPs. The zonation of the diagram is based on unconstrained cluster analysis of palynomorph taxa (see Fig 7).

Fungal remains are common in PZ 2, albeit with wide variations in frequency between barely present and 28%. In PZ 3 and PZ 4 fungal values drop to a few percent (less than 6%) only to rise at the very top of the seam again. Fungal remains do not stray beyond the seam.

This holds true for periderm cells as well. However, contrary to fungal remains periderm cells are nearly absent in the lower part of PZ 2 (PZ 2a) but rise markedly at the base of PZ 2b. With sample S2a-10 they drop back to insignificance only to return, similar to fungal remains, near the top of the seam with maximum values (18%). The marked change in periderm cells from PZ 2a to PZ 2b is accompanied by an equally marked increase of some tricol(po)rate taxa, e.g. *Tricolpopollenites retiformis, T. liblarensis/microhenrici* and *Tricolporopollenites belgicus* (Fig 7A). Cuticle fragments are remnants of leaf cuticles which are easily washed out to the sea and drifting onshore along the shoreline [90, 91]. Accordingly, they appear most frequently in PZ 1 and PZ 5.

Resin (resinite) and tannin-derived bodies (phlobaphinites) are the most common organic components second to charcoal. They represent resistant cell fillings set free from decaying wood and are most abundant in PZ 2a and PZ 2b, but common to frequent throughout the whole seam with considerable fluctuation.

Charcoal particles become the dominant non-palynomorph element in PZ 3 and especially in PZ 4 in striking parallelism to the frequency of *Sphagnum*-type spores and the *Triporopollenites robustus/rhenanus* group. The appearance of pollen of the Normapolles group and freshwater algae (Zygnemataceae) also coincides with the dominance of charcoal in PZ 4.

### Diversity (section S1)

In order to get estimates for palynological richness, rarefaction analyzes of 11 samples from Seam 1 were performed, distinguishing between point diversity within a single sample (Fig 12A, Table 1), alpha diversity within PZs 2 to 4 (Fig 12B, Table 2) and gamma diversity for the entire seam (Fig 12C, Table 2). Furthermore, analysis of beta diversity as well as evenness have been carried out (Tables 1 and 2).

**Fig 12.**
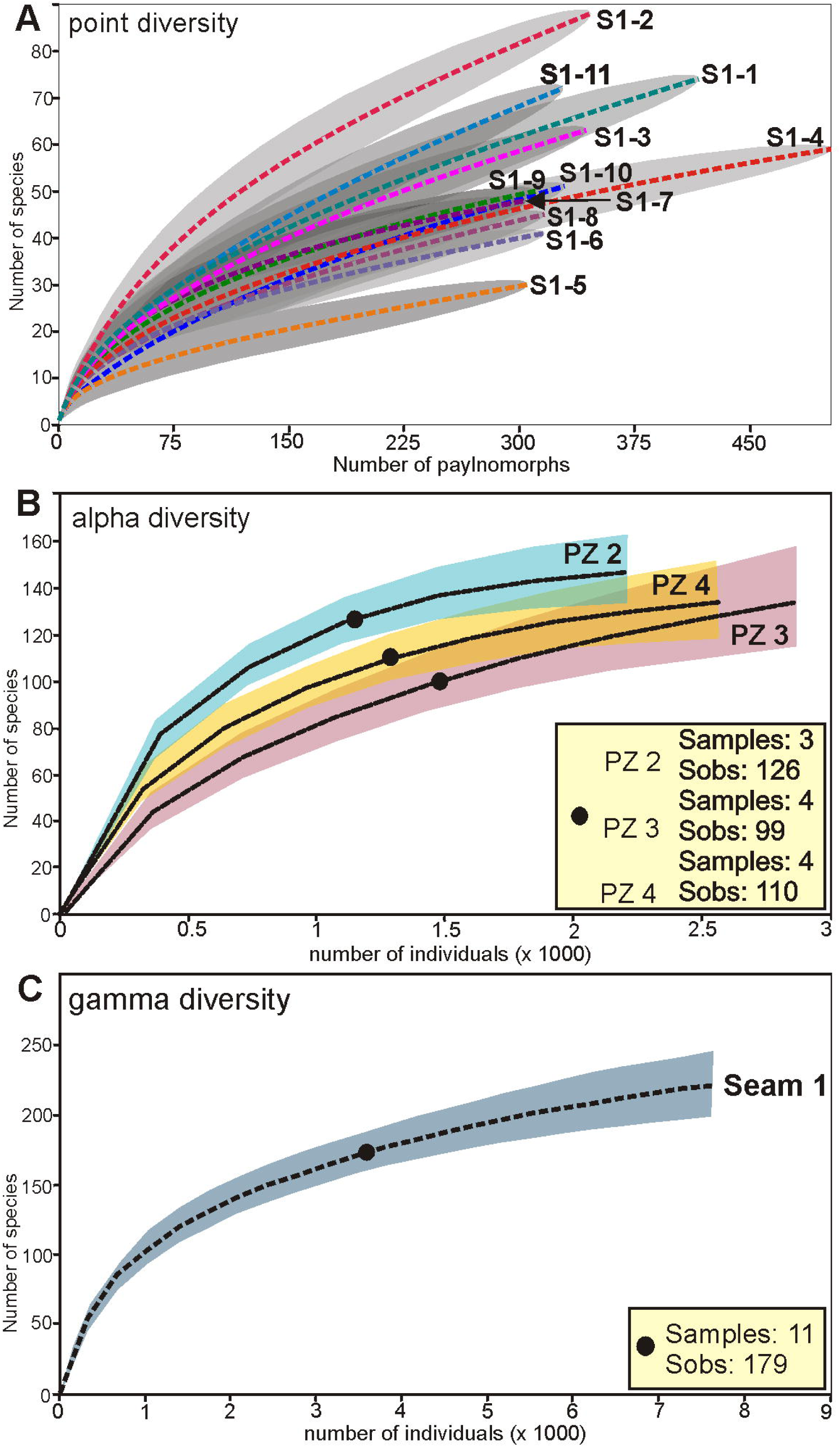
Palynological richness calculations for Seam 1 in section S1 using rarefaction analyses. (A) Point diversity: Individual rarefaction with conditional variance of 11 samples of Seam 1 using the algorithm of [92]. (B) Alpha diversity: Sample-based interpolation and extrapolation using the Bernoulli product model [75] for the 3 palynozones (PZ) of Seam 1 with 95% unconditional confidence intervals; Sobs, number of observed species. (C) Gamma diversity: Sample-based interpolation and extrapolation using the Bernoulli product model [75] for the entire data set of samples from Seam 1; Sobs, number of observed species. Because of differences in the number of counted individuals per sample, the sample-based rarefaction curves and their confidence intervals in (B) and (C) are replotted against an x-axis of individual abundance.

**Table 2:** Estimations of richness and evenness using sample-based incidence data for the 3 palynozones (PZ) of Seam 1 (alpha diversity) and for all samples of Seam 1 (gamma diversity).

|  | PZ 2 | PZ 3 | PZ 4 | Seam 1 |
| --- | --- | --- | --- | --- |
| Number of samples (n) | 3 | 4 | 4 | 11 |
| S<sub>obs</sub> | 126 | 99 | 110 | 179 |
| Individuals counted | 1105 | 1427 | 1283 | 3815 |
| S(est)<sub>5</sub> | 144 | 111 | 119 | 217* |
| S(est)<sub>5</sub> 95% CI, Lower Bound | 131.2 | 96.5 | 107.9 | 196.0* |
| S(est)<sub>5</sub> 95% CI, Upper Bound | 154.7 | 124.6 | 130.5 | 238.3* |
| Singeltons | 49 | 40 | 41 | 58 |
| Doubletons | 21 | 13 | 21 | 20 |
| Shannon-Wiener index | 3.89 | 3.09 | 3.12 | 3.69 |
| Evenness (E) | 0.80 | 0.67 | 0.66 | 0.71 |
| Beta diversity | 0.68 | 1.23 | 1.02 | 2.17 |

S<sub>obs</sub>: Actual number of taxa within the samples; S(est)<sub>5</sub>: Expected number of species in 5 samples using the Bernoulli product model [75]; S(est)<sub>5</sub> 95% CI: Lower and upper bounds of 95% confidence interval for S(est); \*S(est) and \*S(est) 95% CI in 20 samples; Singeltons: Number of species that occur only once in all samples; Doubletons: Number of species that occur only twice in all samples; evenness calculation using the method of [77]: $E = \text{Shannon-Wiener index} / \ln(S_{\text{obs}})$ ; beta diversity using the measure of [78,83]: $(S/\bar{a}) - 1$ (S, total number of species in the PZ or Seam 1; $\bar{a}$ , average number of species in the PZ or Seam 1; \*\* beta diversity estimation without sample S1-5).

Comparison of point diversity within the seam as based on individual rarefaction analyzes using the algorithm of [92], samples of PZ 2 (S1-1 to S1-3) together with sample S1-11 from the top of the seam provides the highest richness values (Fig 12A). While in samples S1-1, S1-3 and S1-11 between 59 and 68 species at 300 counted individuals can be expected, sample S1-2 shows by far the highest number with 82 species (Table 1). The richness in samples from the succeeding PZs 3 and 4 (samples S1-4 to S1-10) is significantly lower with values typically ranging from 40 to 49 species among 300 counted palynomorphs (Table 1). In sample S1-5, the lowest value with only 30 different species is achieved. Therefore, a decrease in palynological richness between PZ 2 and PZs 3 and 4 is significant. Only at the top of the seam in sample S1-11 an increase of richness to values similar to those in PZ 2 is recognizable.

The same pattern of species richness is also evident in alpha diversity (Fig 12B). 126 different pollen and spore taxa have been recorded in the three samples of PZ 2 while significantly lower numbers were observed in the subsequent PZ 3 with 99 species and in PZ 4 with 110 species, although the number of samples and of counted palynomorphs in these two PZs is higher than in PZ 2 (Table 2). Even if the 95% confidence intervals are considered, which describe the range of the possible number of species within the PZs, the richness in PZ 2 is significantly higher than in the two subsequent PZs (Table 2). For PZ 3 and PZ 4 the 95% confidence intervals overlap somewhat (Fig 12B). A diversity increase from PZ 3 to PZ 4 is therefore indicated by the richness estimations, but this is not statistically significant. Furthermore, the interpolation/extrapolation graph of PZ 3 is not saturated indicating that the maximum number of species is higher than calculated and may possibly be in the same range or even higher than in PZ 4.

The high number of singletons and doubletons, showing the number of species with only one or two individuals within the data set, is striking (Table 2). For example, 70 of 126 species of pollen and spores in PZ 2 and 53 of 99 species in PZ 4 have only been recorded one or twice. Therefore, *c*. 55% of the species in the three PZs are singletons or doubletons indicating accordingly that more than half of the species within the total pollen assemblages belong to rare taxa.

The analysis of gamma diversity (Fig 12C) shows a high overall species richness for the entire section. 179 different species have been detected in Seam 1. An extrapolation to 20 samples even indicates a much higher number of morphologically distinct species (217, see Table 2). Since the interpolation/extrapolation graph is not saturated, even more species can be expected (Fig 12C).

Beta diversity as a measure of the difference in species composition is especially high in comparison between sample S1-5 and the other samples with values always higher than 0.6 (S6 Table). This underlines the special composition of the palynomorph assemblage of sample S1-5 in comparison to the other lignite samples of section S1. In contrast, the values for beta diversity of sample comparisons within the same PZs are generally below 0.5 or between 0.5 and 0.6 if samples of different PZs are compared. This indicates minor changes in the composition of the palynomorph assemblages within the PZs, but changes in composition between PZs 2, 3, and 4. This is also confirmed by general beta diversity calculations for the PZs (Table 2). They are low with 0.68 for PZ 2 and 1.02 for PZ 4. Only in PZ 3 the value increases to 1.23, but this is due to the specific composition of the palynomorph assemblage in sample S1-5. If this sample is excluded from the analysis, the value drops to 0.74. In contrast, the total beta diversity value of 2.17 for Seam 1 is significantly higher indicating strong changes in the composition of the palynomorph assemblages between the individual PZs (Table 2).

In addition to species richness, the calculation of evenness provides another important parameter for diversity analysis. In single samples from Seam 1, usually evenness values of more than 0.7 are reached (Table 1). These high values show that the different palynomorph species within the microfloral assemblages are in general evenly distributed in the individual samples. This indicates that (except for the high number of rare elements which contribute to the richness calculation) none of the abundant elements is clearly dominating. Only in samples S1-5 and S1-10 do the evenness values decrease to 0.64 and 0.61 showing that in these samples a dominance of some elements within the pollen assemblage becomes apparent.

PZ 3 and PZ 4 are characterized by relatively low evenness values of 0.67 and 0.66 (Table 2). In contrast, the evenness for PZ 2 is significantly higher with 0.8. Together with the high value for species richness, the high evenness value therefore proves a morpho-diversity in samples of PZ 2 that is significantly higher than in PZ 3 and PZ 4. The evenness value of 0.71 for the entire seam is in accordance with the values of the individual samples (Table 2).

## Discussion

Reconstructions of paleoenvironments from pollen assemblages in peat or coal are mainly based on studies of the relation between standing vegetation and pollen in surface peat samples. Among the numerous studies we prefer to rely on case studies from the coastal plains of the southeastern United States (e.g., [93–95]) which we consider to be rather similar to our example with regard to climate and geologic setting. These studies show that each plant community leaves a distinctive fingerprint in the pollen and spore assemblages of the corresponding peat substrate, although the relationship is not proportional and varies depending f.i. on differences in pollen production, pollination type and preservation potential. The following reconstructions commonly refer to the results of these studies.

### Reconstruction of the paleoenvironment

The NMDS of all three sections show a distinctive threefold succession of vegetation during formation of Seam 1 (Figs 5-7): an initial (PZ 2), a transitional (PZ 3) and a terminal stage (PZ 4). Such tripartite divisions have previously been described and interpreted in terms of environment and vegetation from the Carboniferous of Britain [96–98] and may be a general feature of paralic coals. Mechanisms controlling facies and environment during transgression and regression in peat forming paralic domains have recently been reviewed [99]. Seam 1 is bordered by interbeds I (PZ 1) and II (PZ 5), both showing marine influence and being largely separated from the PZs 2 to 4 in the NMDS of the total data set (Fig 13). Thus in total the following four different types of peat depositional environments and associated vegetation can be distinguished in the three sections: (1) a coastal vegetation (PZ 1 and PZ 5), (2) an initial mire (PZ 2), (3) a transitional mire (PZ 3) and (4) a terminal mire (PZ 4). They remained unaffected by the onset of a warming event at the top of the seam [27] and may therefore be considered as representing plant associations typical for a mid-latitude lowland vegetation during the early Eocene climatic background.

**Fig 13.**
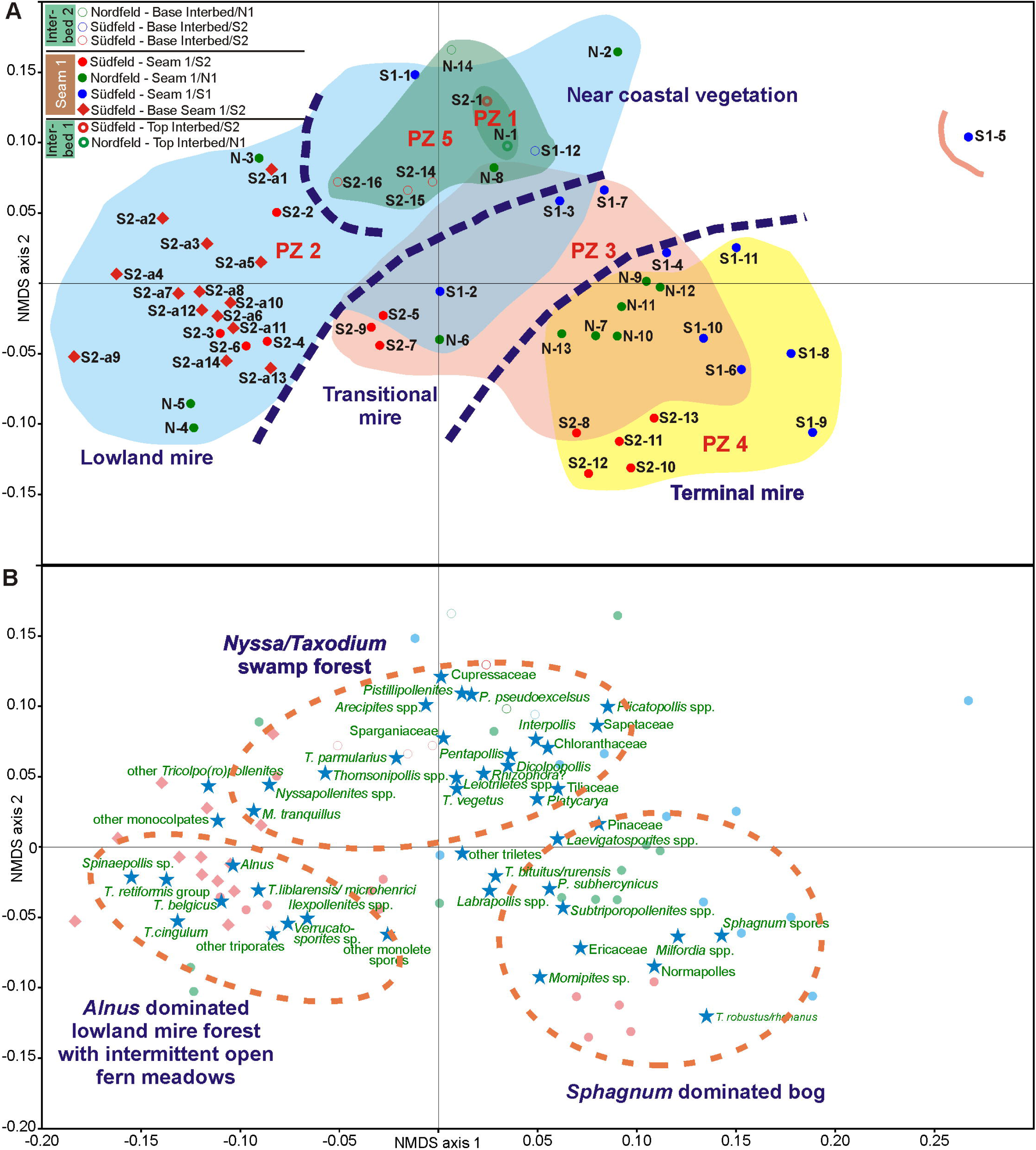
Non-metric multidimensional scaling (NMDS) scatter plots of 56 samples from sections N, S1 and S2. (A) Arrangement of samples (B) Arrangement of palynomorph taxa (stars). The colored dots and diamonds indicate the position of the samples presented in (A). For calculation the Bray-Curtis dissimilarity and Wisconsin double standardized raw-data values have been used.

### Coastal vegetation (PZ 1, PZ 5)

Sandwiched between two marine-influenced interbeds Seam 1 was deposited between a regressive phase represented by PZ 1 (top of Interbed 1) and a transgressive phase represented by PZ 5 (base of Interbed 2). The NMDS of the total data set (Fig 13A) shows that samples of PZ 1 and PZ 5 are largely separated from most of the samples of Seam 1 in the ordination space but plot together with some samples of PZ 2. This indicates that both marine influenced PZs include elements of the peat-forming mire vegetation.

The dominance of *Inaperturopollenites* spp. in PZ 1 and PZ 5 shows that Cupressaceae s.l. played an important role in the coastal vegetation. Together with *Nyssapollenites*, fairly common in the succeeding PZ 2, they indicate that a *Nyssa-Taxodium* swamp forest bordered the coastline at Schöningen and shed the largely wind-driven pollen load into the adjacent estuary. This type of swamp community presently exists in the warm-temperate coastal plains of eastern North America [100, 101]. It has originally been used as a model for the succession of the peat-forming vegetation in the Miocene Lower Rhine Lignite [102, 103] but has later been extended to lignites from other sites of Cenozoic age in Europe [40,104–107]. Since recent *Taxodium*-*Nyssa* swamp forests are common in inland riverine environments [100, 101], their proximity to an estuary at Schöningen appears rather unusual. No particular sedimentological evidence pertaining to a *Nyssa-Taxodium* swamp has been observed in the three sections, but is, however, not expected in a forest derived and intensely rooted peat. Associated elements are *Plicatopollis* spp., *Tricolporopollenites liblarensis* and *Plicapollis pseudoexcelsus*. The latter has been interpreted as a back-mangrove element associated with marsh elements in the middle Eocene Helmstedt Formation [24,108,109]. The anemophilous *Plicatopollis* spp. and *T. liblarensis* as well as the very thin-walled *Inaperturopollenites* spp. are also likely to be derived from nearby external sources such as the background mire forest (Fig 14 A).

**Fig 14.**
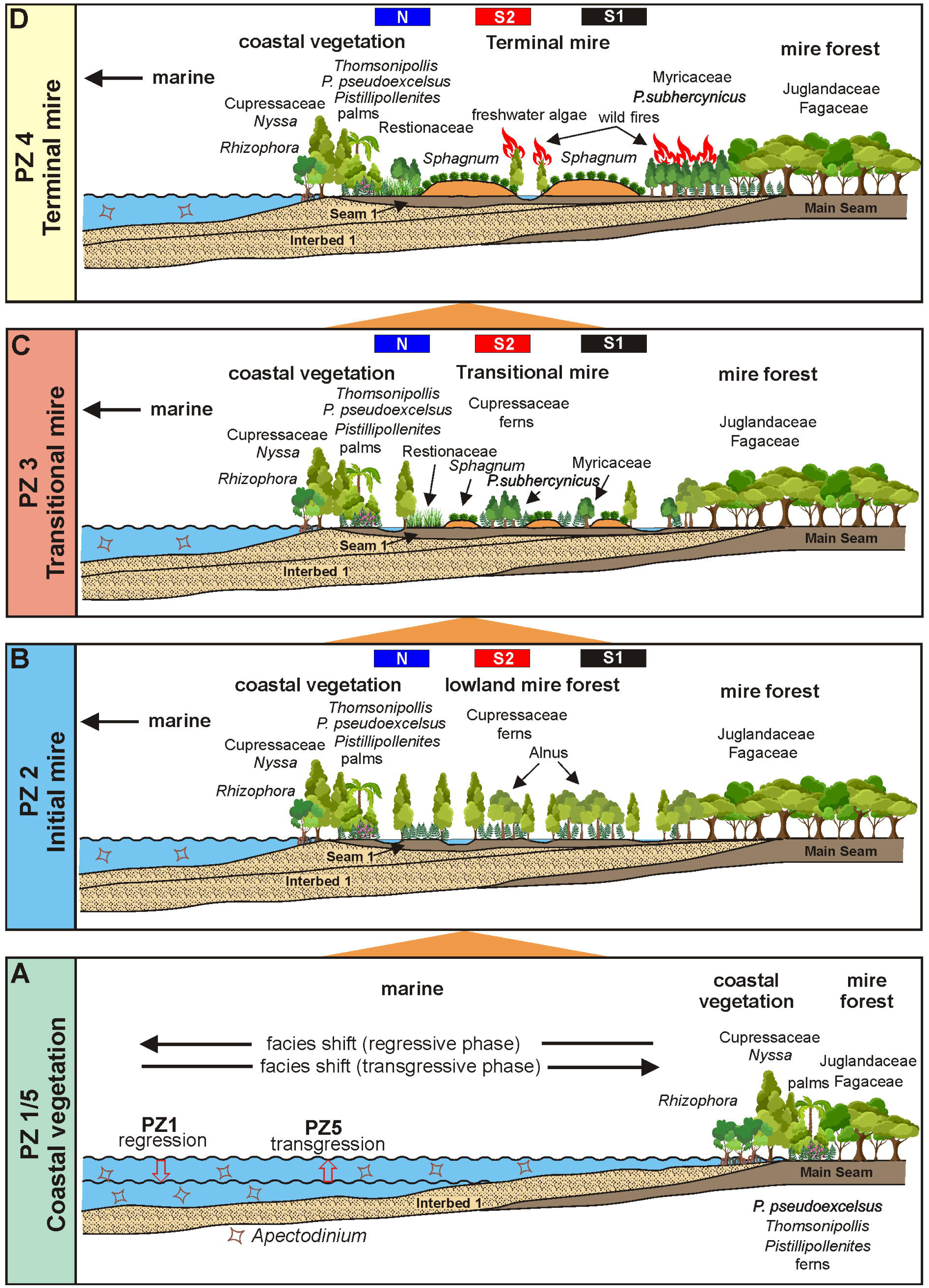
Paleoenvironment reconstruction for Seam 1. Four different types of paleoenvironment and vegetation can be distinguished in the three sections N, S1 and S2. From base to top: (A) a coastal vegetation (PZ 1 and PZ 5) (B) an initial mire (PZ 2) (C) a transitional mire (PZ 3) (D) a terminal mire (PZ 4). The positions of the three studied sections (N, S1, S2) relative to the coast are indicated by colored bars.

Except for scattered occurrences of putative *Rhizophora* (Fig 14A) true mangrove pollen as characterizing the coastal vegetation of the middle Eocene Helmstedt Formation together with *Avicennia* and *Nypa* [24-26,108], is completely missing in the Schöningen Formation [25, 27]. Instead, *Pistillipollenites mcgregorii* and *Thomsonipollis magnificus* (both of unknown botanical affinity) may have substituted here for mangrove elements [25]. Since *T. magnificus* occurs regularly in PZ 1 and 5 in sections S1 and S2 and is very abundant in PZ 2a in section N, where *P. mcgregorii* also occurs at least in low numbers, the parent plants of both taxa were probably common in the immediate coastal vegetation during the deposition of the lower part of the Schöningen Formation.

Finally, cuticle fragments which are abundant in both, PZ 1 and PZ 5, may have had their source in the coastal vegetation and were concentrated along the shoreline by winnowing [90, 91].

### Initial mire (PZ 2)

At the onset of Seam 1 palynomorph assemblages combined in PZ 2 indicate a trend in the vegetation that started in PZ 1 and passes into PZ 3. As shown by the NMDS samples of PZ 2 are plotted together on the negative side of axis 1 in the ordination space (Fig 13A). However, there is little separation from the samples of PZ 1 and PZ 5 and an overlap with samples from the following PZ 3.

The abundance of *Inaperturopollenites* spp. (Cupressaceae s.l.) and the common occurrence of *Nyssapollenites* spp. (Nyssaceae) on either side of the interbed/seam boundary support the existence of a *Nyssa/Taxodium* swamp forest in the immediate vicinity of the coastline. This swamp forest may have been locally replaced by or mixed with patches of other elements, such as the parent plant of *Plicapollis pseudoexcelsus* (base of PZ 2 in section N and S1, Figs 5 and 6), a characteristic element of transitional marine/terrestrial environments of possible juglandaceous affinity [24,108,109]. *Thomsonipollenites magnificus* is quite abundant in section N (PZ 2a) in contrast to the other two sections.

In particular, *Alnipollenites verus* (*Alnus*) is common to frequent throughout PZ 2 and even highly dominant in some samples of section N (e.g., N4, Fig 5) and S2 (e.g., S2-a2, Fig 7). For these sites temporarily inundated freshwater wetland habitats may be envisioned similar to those in which modern species of *Alnus* such as e.g. *A*. *glutinosa* [110], *A*. *incana* [111] or *A*. *viridis* [112] grow today. Intermittent open fern meadows are indicated by the strong proliferation of trilete spores at the base of PZ 2 in section N. Notably, these spores are absent in the other two sections. Other common associates of PZ 2 assemblages such as *Monocolpopollenites tranquillus* (Arecaceae, *Phoenix*), *Plicatopollis* spp., and *Tricolpopollenites liblarensis* may have been in part indigenous to PZ 2, but they are small and thin-walled and therefore considered to be anemophilous [41] and likely to be introduced from other sources.

Local differences shown in the three sections are a special feature of PZ 2 indicating a pronounced patchiness in the initial mire vegetation (Fig 14B). This may be due to a number of variables controlling the structure of mire vegetation such as minute relief in the sub-peat topography [113, 114], the water quality (salinity, nutrient load) or different dispersal strategies and competition among plants or other localized disturbances interrupting directional successions [115] which together result in differential peat aggradation [116].

This is also reflected in the striking contrast between subzones PZ 2a and PZ 2b in sections N and S2. Notable is, for instance, the replacement of fern spores (*Leiotriletes* spp.) in section S2 by pollen of woody plants such as *Tricolporopollenites belgicus* (Fig 7). The change from a herbaceous vegetation rich in ferns in PZ 2a to a more woody vegetation in PZ 2b is even reflected in the distribution of non-palynomorph organic remains showing an increase of periderm cells, phlobaphinites and resin particles as well as fungal remains from PZ 2a to PZ 2b (Fig 11).

### Transitional mire (PZ 3)

The change in vegetation occurring within PZ 3 is gradual. In the course of peat aggradation previously dominant elements such as *Alnipollenites verus* (*Alnus*) or *Inaperturopollenites* spp. (Cupressaceae s.l.) are replaced by taxa such as *Pompeckjoidaepollenites subhercynicus* as well as *Sphagnum*-type spores. The latter two become eventually dominant in the succeeding PZ 4 thus indicating the transitional character of this pollen zone and a reduction of (temporally) flooded habitats.

Accordingly, the samples of PZ 3 plot midway between those of the clearly separated PZ 2 and PZ 4 in the ordination space of the NMDS of the total data set (Fig 13A). There are, however, considerable areas of overlap with both PZs which characterize PZ 3 as transitional between the initial and the terminal phases in the formation of Seam 1. The PZ 3 samples of section S2 differ from those of the other two sections since they plot separate to the left on the negative side of axis 1 (Fig 13A). This is due to the fact that the similarity of samples from S2 to those of PZ 2 is closer than in the other two sections, which are more transitional to PZ 4.

*P. subhercynicus* (unknown botanical affinity) and the *T. robustus/rhenanus* group pollen (Myricaceae) are widely distributed throughout the Schöningen Formation and are often dominant in the upper part of the lower seams (Main Seam, Seam 1 and Seam 2 [27]). *P. subhercynicus* is more restricted to certain levels and appears to prefer mire forest/marsh interfaces (ecotones) [24, 26]. The *T. robustus/rhenanus* group is locally abundant in PZ 3 and even dominant in section N before becoming dominant throughout PZ 4. Noteworthy are the first peaks of *Sphagnum*-type spores indicating an initial tendency for ombrogenous bogs to develop under the open canopy of an angiosperm mire forest (Fig 14C).

Sample 5 of section S1 is clearly separated from all other samples in this section in the NMDS (Figs 6C, 13) by the dominance of peat moss and fern spores (*Sphagnum*-type spores, *Laevigatosporites* spp*., Leiotriletes* spp.) together with a mass occurrence of charcoal particles. The sample was taken from a layer (X-Horizon), which included tree-stumps as well as a charcoal horizon (Figs 4 and 6). Possibly, tree fall and/or forest fire may have left a clearing here, which was resettled by ferns and mosses as pioneers [117].

### Terminal mire (PZ 4)

A very marked change in palynomorph assemblage composition occurs at the transition from PZ 3 to PZ 4. This is mainly due to the rise to dominance of *Sphagnum*-type spores including all three genera previously observed in seams of the Schöningen Formation, i.e. *Sphagnumsporites, Tripunctisporis* and *Distancorisporis* [44, 89]. Although the latter two are morphologically different from modern *Sphagnum* spores the three genera are sufficiently similar and closely associated with remains of *Sphagnum* (leaves) in a thin lignite seam (*Sphagnum* Seam) higher up in the Schöningen section to confirm their affinity to *Sphagnum* [25,44,89].

The change in PZ 4 is underscored by the great increase in pollen of the *T. robustus/rhenanus* group. Although some of these changes are already initiated in section N (for *T. robustus/rhenanus*) and section S1 (for *Sphagnum-*type spores), the NMDS of all three sections (Figs 5C, 6C, 7C) and of the total data set (Fig 13A) show a clear separation of PZ 4 samples from those of all other PZs.

A number of authors have affiliated *T. robustus* with various, mostly catkin-bearing families. But on the basis of surface features visible at high resolution ([118, 119], personal SEM observation) we favour an affinity with the Myricaceae, a family today represented mostly by small trees and shrubs adapted to wet acidic environments and nutrient deficiency [120, 121]. Together with *Sphagnum* they clearly signal that peatbeds in PZ 4 were decoupled from groundwater (raised bog) and their hydrology increasingly, but not exclusively controlled by precipitation since freshwater runoff was backed up by the rise of sea level [122–124] (Fig 14D). The increase of *Ericipites* sp. (Ericaceae) and *Milfordia* spp. (Restionaceae) in section S1 is fully in line with this development. In particular, Restionaceae have been described as an important constituent of the so-called *Sphagnum* Seam at Schöningen which has been compared with recent southern hemisphere restionad bogs [44]. Pools of standing water are a common feature of terminal mires [125–127] and indicated here by the rare, but regular occurrence of remains of freshwater algae such as cysts of Zygnemataceae (Fig 14D).

Somewhat intriguing are certain members of the Normapolles such as *Basopollis* and *Nudopollis*, relics from the Cretaceous, the occurrence of which is one of the last in the Paleogene of Central Europe and largely restricted here to PZ 4. Their parent plants seem to have found refuge within the vegetation and environment of PZ 4 just prior to their extinction. The association of *Basopollis orthobasalis*, f.i., with pollen of Myricaceae indicates that the parent plants of Normapolles favored the nutrient deficient conditions of the terminal mire of PZ 4 [128].

The multiple evidence of waterlogged conditions and standing water, however, seems counterintuitive to the massive occurrence of charcoal particles (Fig 11). Since they often show bordered pits, they are mostly wood derived and, thus, clearly differing from the charcoal of the herb-dominated early Eocene *Sphagnum* bog at Schöningen [44]. This apparent contradiction may be resolved in three ways: by close lateral proximity of burnt and waterlogged to aquatic sites, by crown fires in a temporarily flooded mire forest or by periodic drought followed by flooding and resettlement of burned forest sites. As a possible modern equivalent for the latter we consider the complex fire regime in the Okefenokee Swamp (Georgia, USA) [129], where periodic forest fires at approximately 25 year intervals left charcoal horizons and lowered the peat surface thus creating space for ponds or lakes when water level returned to normal [130, 131]. New peat was deposited after each fire [132]. In PZ 4 of Seam 1 new peat was formed among others by regrowth of *Sphagnum*, ferns, Restionaceae (section S1), and shrubs (Myricaceae, Betulaceae, Juglandaceae).

### Comparison of palynomorph assemblages between sections N, S1, S2

The abundance values of common taxa are averaged for the five PŹs in each of the three sections and juxtaposed in Fig 15, in order to show the differences between sections as opposed to the vertical succession of pollen assemblages. A gradient from the most seaward (section N) to the furthest inland section (section S2) becomes evident for several taxa, which appear to be environmentally sensitive.

**Fig 15.**
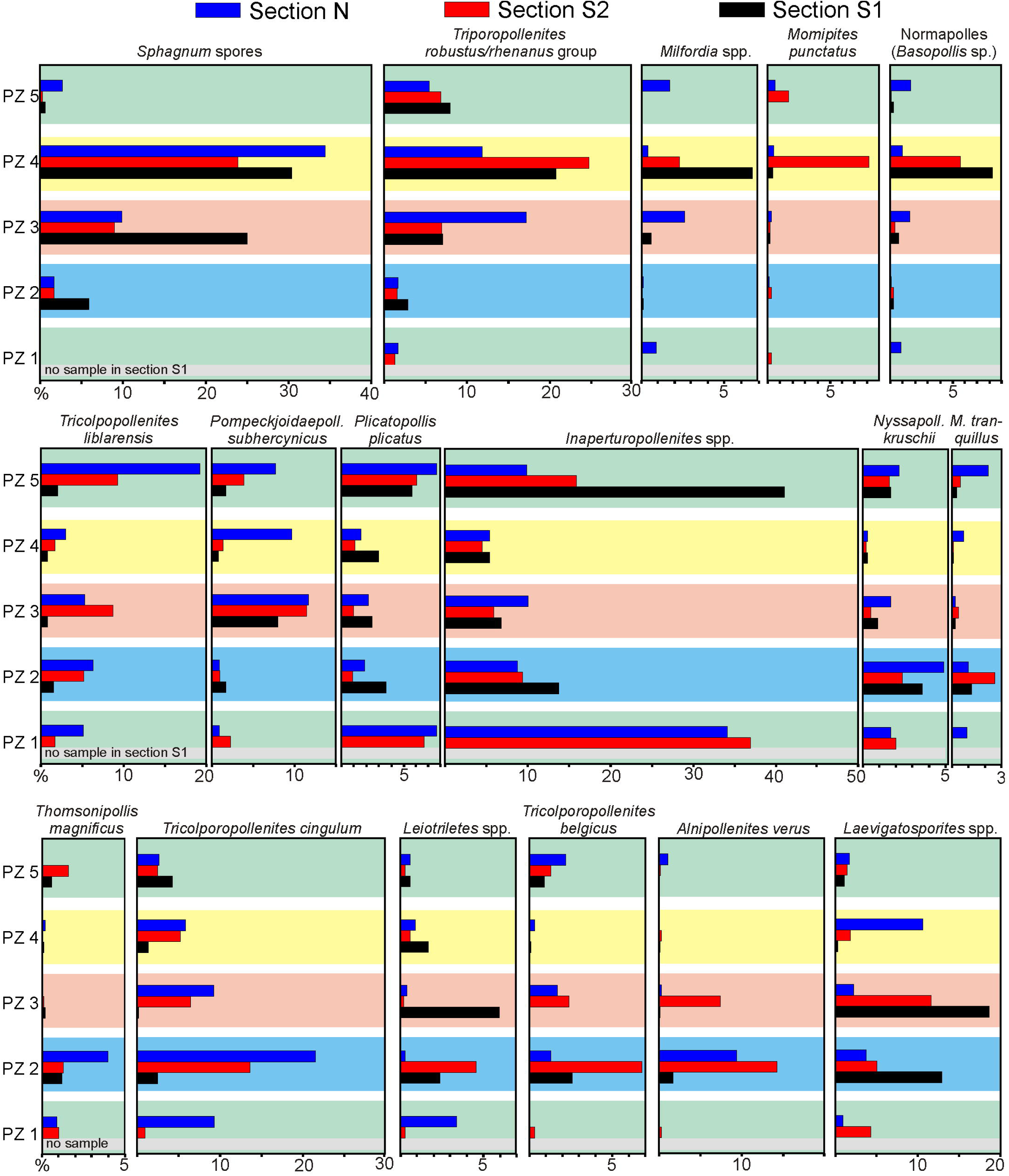
Average abundance of important palynomorph taxa in the different palynozones of the three sections of Seam 1. Average values are presented in percent for each of the PZs. The three sections are arranged from top to bottom away from the coastline.

Thus, taxa, which have previously been found in association with marine influence, are best represented in section N. *Thomsonipollis magnificus*, for example, has been considered to substitute for the true mangrove elements in various sections at Helmstedt and Schöningen, while *Pompeckjoidaepollenites subhercynicus* appears to be an important constituent of the back-mangrove ecotone [24,25,108]. *Nyssa* (*Nyssapollenites kruschii*) and *Taxodium* (*Inaperturopollenites* sp. partim) are most tolerant of standing freshwater among trees and, thus, are likely to have formed stands in adjacent backwater areas. The small tricolp(or)ate taxa (*Tricolpopollenites liblarensis, Tricolporopollenites cingulum*) may be considered to be wind-driven from more inland mire sites and proportionately overrepresented as other elements decline closer to the shore.

Corresponding to the successional upward increase there is also a landward increase in the abundance of Sphagnum-type spores indicating the initiation of Sphagnum peat bogs in more inland sites. Moreover, the diagram (Fig 15) indicates that they may have been first developed in the most landward site, where they occur already in PZ 3. A similar trend of abundance increases towards more inland sites is shown by pollen of Myricaceae (*T. robustus/rhenanus* group), Restionaceae (*Milfordia* spp.) and Normapolles (*Basopollis*, unknown botanical affinity), in part also for Juglandaceae (*Momipites punctatus*).

Fern spores (*Leiotriletes, Laevigatosporites*) as well as *Alnipollenites* (*Alnus*) and to some extent *Tricolpopollenite belgicus* show no clear pattern and are characterized by strong frequency fluctuations in the three sections, which agrees well with their potential role as invaders of locally disturbed sites.

Overall, it should be noted that, despite the differences between the three sections, differences in the quantitative composition of pollen assemblages are significantly more pronounced between successional stages as represented by the pollen zones. Thus, the five pollen zones distinguished here are a robust fingerprint of the successional stages of vegetation, under which Seam 1 has been formed.

### Diversity

The study of the morpho-diversity of the pollen assemblages in section S1 allows for an estimate of the diversity of the vegetation, assuming that the pollen rain reflects relative changes within the vegetation [77, 80]. 179 palynomorph species have been recognized in Seam 1, but gamma diversity calculation shows that more than 200 species can be expected (Fig 12C, Table 2). Thus, the diversity is higher than reported for other lower Eocene records, especially those in Central Europe such as Krappfeld (Carinthia, Austria) [133, 134], Epinois (Belgium) [56] or the Cobham Lignite (southern England) [22] and calculated for North America (Mississippi, Alabama) [135, 136]. However, the calculations may be based on different taxonomic resolution and should therefore be considered with caution. In any case, the diversity measures of Seam 1 reflect a high plant diversity as typical for forested tropical coastal wetlands and peatlands [137].

Point and alpha diversities of PZ 2 in section S1 are significantly higher than in the other PZs in S1 (Figs 12A and 12B). This high morpho-diversity of the microflora in PZ 2 is rather striking, but not surprising when PZ 2 is considered to represent an ecotone between PZ 1 and PZ 3 combining elements from the coastal vegetation and the subsequent initial mire forest. Furthermore, an initial mire forest, as represented in PZ 2 is in any case highly diverse compared to a disturbed terminal mire vegetation with raised bog that follows later in PZ 4 [137]. With the initiation of a *Sphagnum* bog in PZ 4 a peat swamp developed that became increasingly oligotrophic supporting plant communities that are adapted to low pH and nutrient depletion and which are low in diversity [138]. Accordingly alpha diversities of PZs 3 and 4 are significantly lower than in PZ 2 (Fig 12B).

Point diversity increases again in the uppermost sample of PZ 4 (Fig 12A) immediately prior to the transgression of Interbed 2, where species-rich back swamp and coastal communities returned to the site. The exceptionally low point diversity and evenness of sample S1-5 (X-horizon) is due to the dominance of spores and the concomitant decline of other elements (Fig 12A, Table 1) possibly caused by resettlement of a clearing in the mire forest by ferns and mosses.

### Paleoclimate

Isotope analyses have recently shown that a Carbon Isotope Excursion (CIE) indicates a short-term thermal event that started with the topmost sample of Seam 1 extending into the lower part of Seam 2 [27]. Nevertheless, the bulk of Seam 1 was deposited during a moderately warm period of the lower Eocene as suggested below. However, temperature reconstructions for Seam 1 based on biomarker analysis (brGDGTs) resulted in high mean annual temperatures (MAT), which reached 24°C ± 4.6°C in the lower part of the seam [139]. Therefore, a thermophilic vegetation should be expected similar to other sites along the southern coast of the Proto-North Sea such as Cobham (southern England) and Vasterival (France) which included the PETM [22, 23].

We present evidence here that the vegetation of Seam 1 indicates a cooler mesothermal climate. True humid tropical mangrove elements such as *Avicennia* and *Nypa*, common in the coastal vegetation of the succeeding middle Eocene Helmstedt Formation [24, 108], are absent. This suggests at least extratropical conditions for the Schöningen Formation [25]. On the other hand, *Alnus*, one of the characteristic elements of PZ 2 does not occur in the PETM records of the Cobham lignite and Vasterival [22, 23]. The assemblages of PZ 2 are more compatible with high-latitude Eocene swamp forests such as those on Axel Heidberg Island in the Canadian High Arctic, where Cupressaceae s.l. and *Alnus* are widely distributed [140]. A similar microflora is also known from the Paleocene/Eocene boundary in the central North Sea [141], where the vegetation is composed of a mixture of azonal and zonal elements including mesothermal conifers (Cupressaceae s.l.) and dicots such as *Alnus*, *Carya* and *Juglans* indicating a mixed conifer broadleaf vegetation [141]. Temperature reconstructions for this record based on comparisons with nearest living relatives (NLR) indicate relatively cool mean annual temperatures (MAT) of 15° C and cold month mean temperatures (CMMT) of 8° C but warm month mean temperatures (WMMT) of 22.5° C for the North Sea region [141]. The similarly composed mixed palynomorph assemblages of PZ 2, in particular the high abundance of *Alnus* pollen, would, therefore, suggest similar extratropical conditions for Seam 1. This is a considerably cooler estimate than that based on biomarker analysis notwithstanding the resemblance of WMMT estimates. However, this may be explained by the fact that a certain temperature bias between brGDGTs estimates and those from leaves and palynomorphs is well known [141, 142].

Our results are consistent with data for the Bighorn Basin in North America where the thermal event of the PETM is followed by a strong temperature decline in the first million years of the Eocene, shortly before the rapid increase of temperatures during the succeeding EECO followed [143]. Seam 1 has been deposited in the lowermost Eocene during a phase following the PETM and before a lower Eocene thermal event in the succeeding Interbed 2 and Seam 2 appeared [27]. Thus, a simultaneous cooling in the continental climate of North America and Central Europe is indicated.

Although *Alnus* as a temperate climate element declines in PZ 2 and PZ 3 extratropical conditions seem to have persisted through PZ 3 and PZ 4 since *Sphagnum* and fern spores in association with pollen of Restionaceae and Ericaceae dominate [25, 44]. They are typical for temperate mires in the southern hemisphere today [144–149]. In the northern hemisphere similar pollen assemblages are also known from Paleocene to lower Eocene coals of Texas and Wyoming [150].

The close association of *Sphagnum* and fern spores with high abundances of charcoal in PZ 3 and PZ 4 (Fig 11) appears rather contradictory and has been interpreted in a number of ways. In any case, the great increase in charcoal points to an increase in fire activity and possibly to dryer conditions in the area toward the end of Seam 1 formation. In a recent study of charcoals from 11 autochthonous lignite seams from the Schöningen and Helmstedt formations, including Seam 1, a mix of charcoal particle sizes from >500 μm to less than 10 μm has been recognized [28], which indicates local and regional wildfire activity [151–154]. Especially larger charcoal particles (≥500 μm) in the autochthonous early Paleogene lignites at Schöningen, such as Seam 1, are unlikely to have been washed in and therefore represent locally occurring wildfires [28]. Semifusinite and inertodetrinite are the most common inertinite macerals in the lignite. Since inertodetrinite is predominantly wind-blown due to its small size [155], it can be used as an indicator of local and regional high temperature crown fire activity [154–156].

Inglis et al. [89] argued that wildfires impeded the spread of taller and more vulnerable vascular plants and thereby advanced the spread of *Sphagnum*. On the other hand, the highest abundance of charcoal at the top of PZ 4 (Fig 11) may be correlated with the onset of a CIE [27] considering that an increase of wildfires shortly before the onset of the PETM has been noted for the Cobham lignite [22, 157]. This could give support to the suggestion that peat burning may have been a trigger for CIEs and associated thermal events in the early Paleogene [158, 159]. However, we favor the Okefenokee Swamp (Georgia, USA) as a recent example for conditions existing during PZ 4, in which periodic droughts and subsequent forest fires leave open areas later invaded by a herbaceous vegetation consisting of *Sphagnum*, ferns, Restionaceae and Ericaceae with aquatic sites in between [160].

## Conclusions

Statistical scrutiny by means of Cluster analyses and NMDS shows that 5 different PZs occurring in vertical succession can be clearly distinguished in the three sections of Seam 1 despite local differences between them. They reflect vegetation responses to changes in environment and facies that took place during an early Paleogene regression/transgression cycle including the formation of a coal seam. The two PZs bounding the seam, PZ 1 and PZ 5, are similar mainly due to the presence of marine indicators (*Apectodinium*, *Rhizophora*) and reflect the state of vegetation during the regressional respectively transgressional phase. PZ 2 to PZ 4 reflect changes occurring during coal seam respectively peat bed formation, in which the various mire communities competed for changing hydrologic conditions, nutrient resources and effects of peat aggradation. The initial phase (PZ 2) was characterized by a patchy, pioneering vegetation (e.g. *Thomsonipollis magnificus*, *Alnipollenites verus*) controlled by variable edaphic conditions. PZ 3 appears transitional in a seam of limited thickness, but represents a certain climax in mire development since it is composed of a mix of species adapted to these conditions. External factors such as fire, flooding and an increasing influence of precipitation led to environmental disturbances and differential peat aggradation supporting a rather heterogeneous vegetation of *Sphagnum*, ferns, and Myricaceae in combination with frequent charcoal (PZ 4) during the terminal phase. A renewed transgression finally truncated further peat aggradation preventing full development of ombrogeneous conditions which commonly constitute the bulk of thick seams as described for brown coals of Victoria, Australia [107].

Diversity measurements show that PZ 2 has the greatest species diversity as is commonly the case in ecotones containing elements from adjacent communities as well as specialists of different habitats. Since they disappear with progressive stabilization of the mire environment, diversity drops to the lowest in PZ 3, before disturbances in the environment create new habitats in PZ 4. This pattern may be considered typical of vegetation responses in regression/transgression cycles.

Climatic signals for Seam 1 are somewhat contradictory. Warm temperatures of *c.* 24 °C have been calculated by biomarker analyses of Seam 1 approaching those accepted for the PETM [139]. Isotope analyses [27], on the other hand, have shown that Seam 1 has been formed just prior to a negative CIE excursion. There is strong palynological evidence from Seam 1 that a temperate climate prevailed in northwestern Germany during the lowermost lower Eocene, since *Alnus* and *Sphagnum* are abundant temperate elements in Seam 1, while tropical elements, e.g. *Avicennia*, *Nypa* and *Sapotaceae*, well known from the middle Eocene Helmstedt Formation, are entirely missing. Seam 1, therefore, stands as an example typical for the normal climate during the early Eocene. As such it will serve as a standard by which vegetation responses to any of the known early Eocene thermal events can be identified in the Schöningen section.

## Author contributions

Conceptualization: Olaf K. Lenz, Walter Riegel, Volker Wilde

Data Curation: Olaf K. Lenz

Formal Analysis: Olaf K. Lenz

Funding Acquisition: Olaf K. Lenz

Investigation: Olaf K. Lenz, Walter Riegel, Volker Wilde

Methodology: Olaf K. Lenz, Walter Riegel, Volker Wilde

Project Administration: Olaf K. Lenz, Volker Wilde

Resources: Olaf K. Lenz, Walter Riegel, Volker Wilde

Validation: Olaf K. Lenz

Visualization: Olaf K. Lenz

Writing – Original Draft Preparation: Olaf K. Lenz

Writing – Review & Editing: Walter Riegel, Volker Wilde

## Acknowledgements and funding

The authors thank Karin Schmidt for valuable field support and technical assistance. We are also grateful to the Helmstedt Revier GmbH (formerly BKB and later EoN) for access to the sections and technical support in the field. We thank 2 anonymous referees for their reviews and comments that helped to improve the manuscript significantly.

## Supporting information

**S1 Table. Taxonomic list**.

Complete list of palynomorphs from the studied sections N, S1, S2 from Seam 1 of the Schöningen Formation including their systematic affinities. In the left column the 45 “variables” are presented, which were used for the pollen diagrams and statistical analysis (cluster analysis, non-metric multidimensional scaling).

**S2 Table: Raw data set of section N.**

The data have been used for pollen diagram, cluster analysis and NMDS.

**S3 Table: Raw data set of section S1 (a).**

The data have been used for pollen diagram, cluster analysis and NMDS.

**S4 Table: Raw data set of section S2**

The data have been used for pollen diagram, cluster analysis and NMDS.

**S5 Table: Raw data set of section S1 (b)**

The data have been used for diversity analysis.

**S6 Table: Estimations of beta diversity for Seam 1 in section S1.**

Given are pairwise comparisons of 11 lignite samples from section S1 using the measure of [59, 64]: (S/ā) – 1; S, total number of species in the two compared samples, ā, average number of species in the two compared samples of Seam 1.

## References

1. Zachos JC, Pagani M, Sloan L, Thomas E, Billups K. Trends, rhythms, and aberrations in global climate 65 Ma to present. Science 2001; 292: 686–693.

2. Kennett JP, Stott LD. Abrupt deep-sea warming, palaeoceanographic changes and benthic extinctions at the end of the Palaeocene. Nature 1991; 353: 225–229.

3. Bains S, Norris RD, Corfield RM, Faul KL. Termination of global warmth at the Palaeocene/Eocene boundary through productivity feedback. Nature 2000; 407: 171–174.

4. Röhl U, Bralower TJ, Norris RD, Wefer G. New chronology for the late Paleocene thermal maximum and its environmental implications. Geology 2000; 28: 927–930.

5. Röhl U, Westerhold T, Bralower TJ, Zachos JC. On the duration of the Paleocene-Eocene thermal maximum (PETM). Geochemistry, Geophysics, Geosystems 2007; 8: Q12002. doi:10.1029/2007GC001784, 2007.

6. Westerhold T, Röhl U, Frederichs T, Agnini C, Raffi I, Zachos JC, et al. Astronomical calibration of the Ypresian timescale: Implications for seafloor spreading rates and the chaotic behaviour of the solar system? Climate of the Past 2017; 13: 1129–1152.

7. Zeebe RE, Lourens LJ. Solar System chaos and the Paleocene–Eocene boundary age constrained by geology and astronomy. Science 2019; 365: 926–929.

8. Lourens LJ, Sluijs A, Kroon D, Zachos JC, Thomas E, Röhl U, et al. Astronomical pacing of late Palaeocene to early Eocene global warming events. Nature 2005; 235: 1083–1087.

9. Sluijs A, Schouten S, Donders T., Schoon PL, Röhl U, Reichart GJ, et al. Warm and wet conditions in the Arctic region during Eocene Thermal Maximum 2. Nature Geosciences 2009; 2: 777–780.

10. Cramer BS, Wright JD, Kent DV, Aubry MP. Orbital climate forcing of δ in the late Paleocene–early Eocene (chrons C24n-C25n). Paleoceanography 2003; 18: 1097. doi: 10.1029/2003PA000909.

11. 11. Röhl U, Westerhold T, Monechi S, Thomas E, Zachos JC, Donner B. The Third and Final Early Eocene Thermal Maximum: Characteristics, Timing and Mechanisms of the “X” Event. Geological Society of America, Abstracts with Programs. 2005; 37: 264.

12. Zachos JC, Wara MW, Bohaty S, Delaney ML, Petrizzo MR, Brill, A., et al. A transient rise in tropical sea surface temperature during the Paleocene-Eocene thermal maximum. Science 2003; 302: 1551–1554.

13. Zachos JC, Dickens GR, Zeebe RE. An early Cenozoic perspective on greenhouse warming and carbon-cycle dynamics. Nature 2008; 451: 279–283.

14. Zachos JC, McCarren H, Murphy B, Röhl U, Westerhold T. Tempo and scale of late Paleocene and early Eocene carbon isotope cycles: Implications for the origin of hyperthermals. Earth and Planetary Science Letters 2010; 299: 242–249.

15. Sluijs A, Dickens GR. Assessing offsets between the δ and the global exogenic carbon pool across early Paleogene carbon cycle perturbations. Global Biogeochemical Cycles 2012; 26. GB4005, https://doi.org/10.1029/2011GB004224.

16. Dickens GR. Carbon addition and removal during the Late Palaeocene Thermal Maximum: Basic theory with a preliminary treatment of the isotope record at ODP Site 1051, Blake Nose. Geological Society London Special Publications 2001; 183: 293–305.

17. Turner SK, Sexton P, Charles CD, Norris RD. Persistence of carbon release events through the peak of early Eocene global warmth. Nature Geosciences 2014; 7: 748–751.

18. Clyde WC, Gingerich PD. Mammalian community response to the latest Paleocene thermal maximum: An isotaphonomic study in the northern Bighorn Basin, Wyoming. Geology 1998; 26: 1011–1014.

19. Bowen GJ, Clyde WC, Koch PL, Ting S, Alroy J, Tsubamoto T, et al. Mammalian dispersal at the Paleocene/Eocene boundary. Science 2002; 295: 2062–2065.

20. Jaramillo C, Ochoa D, Contreras L, Pagani M, Carvajal-Ortiz H, Pratt LM, et al. Effects of rapid global warming at the Paleocene-Eocene Boundary on Neotropical Vegetation. Science 2010; 330: 957–961.

21. Wing SL, Currano ED. Plant response to a global greenhouse event 56 million years ago. American Journal of Botany 2013; 100: 1234–1254.

22. Collinson ME, Steart DC, Harrington GJ, Hooker JJ, Scott AC, Allen LO, et al. Palynological evidence of vegetation dynamics in response to palaeoenvironmental change across the onset of the Paleocene-Eocene Thermal Maximum at Cobham, Southern England. Grana 2009; 48: 38–66.

23. Garel S, Schnyder J, Jacob J, Dupuis C, Boussafir M, Le Milbeau C, et al. Paleohydrological and paleoenvironmental changes recorded in terrestrial sediments of the Paleocene–Eocene boundary (Normandy, France). Palaeogeography, Palaeoclimatology, Palaeoecology 2013; 376: 184–199.

24. Lenz OK. Palynologie und Paläoökologie eines Küstenmoores aus dem Mittleren Eozän Mitteleuropas-Die Wulfersdorfer Flözgruppe aus dem Tagebau Helmstedt, Niedersachsen, Palaeontographica B 2005; 271: 1–157.

25. Riegel W, Wilde V, Lenz OK. The Early Eocene of Schöningen (N-Germany) – an interim report. Austrian Journal of Earth Sciences 2012; 105: 88–109.

26. Riegel W, Lenz OK, Wilde V. From open estuary to meandering river in a greenhouse world – An ecological case study from the Middle Eocene of Helmstedt, northern Germany. Palaios 2015; 30: 304–326.

27. Methner K, Lenz OK, Riegel W, Wilde V, Mulch A. Paleoenvironmental response of midlatitudinal wetlands to Paleocene–early Eocene climate change (Schöningen lignite deposits, Germany). Climate of the Past 2019; 15: 1741–1755.

28. Robson BE, Collinson ME, Riegel W, Wilde V, Scott AC, Pancost RD. Early Paleogene wildfires in peat-forming environments at Schöningen, Germany. Palaeogeography, Palaeoclimatology, Palaeoecology 2015; 437: 53–62.

29. Brandes C, Pollok L, Schmidt C, Wilde V, Winsemann J. Basin modelling of a lignite-bearing salt rim syncline: insights into rim syncline evolution and salt diapirism in NW Germany. Basin Research 2012; 24: doi: 10.1111/j.1365-2117.2012.00544x.

30. Blumenstengel H, Krutzsch, W. Tertiär. In: Bachmann GH, Ehling BC, Eichner R, Schwab M, editors. Geologie von Sachsen-Anhalt. Stuttgart: Schweizerbart; 2008. pp. 267– 273.

31. Standke G. Paläogeografie des älteren Tertiärs (Paleozän bis Untermiozän) im mitteldeutschen Raum. Zeitschrift der Deutschen Gesellschaft für Geowissenschaften 2008; 159: 81–103.

32. Wilde V, Riegel W, Lenz OK. Das Paläogen im Helmstedter Revier: Ein Forschungsthema im Geopark Harz. Braunschweiger Land. Ostfalen. Gaussiana 2020; 1: Forthcoming.

33. Ziegler PA. Geological Atlas of Western and Central Europe. The Hague: Shell Internationale Petroleum Maatschappij B.V; 1990.

34. Manger G. Der Zusammenhang von Salztektonik und Braunkohlenbildung bei der Entstehung der Helmstedter Braunkohlenlagerstätten. Mitteilungen aus dem Geologischen Staatsinstitut in Hamburg 1952; 21: 7–45.

35. Baldschuhn R, Binot F, Fleig S, Kockel F. Geotektonischer Atlas von Nordwest-Deutschland und dem deutschen Nordsee-Sektor. Geologisches Jahrbuch Reihe A 1996; 153: 1–88.

36. Gramann F, Harre W, Kreuzer H, Look ER, Mattiat B. K-Ar ages of Eocene to Oligocene glauconitic sands from Helmstedt and Lehrte (Northwestern Germany). Newsletter on Stratigraphy 1975; 4: 71–86.

37. Gürs K. Das Tertiär Nordwestdeutschlands in der Stratigraphischen Tabelle von Deutschland 2002. Newsletters on Stratigraphy 2005; 41: 313–322.

38. Ahrendt H, Köthe A, Lietzow A, Marheine D, Ritzkowski S. Lithostratigraphie, Biostratigraphie und radiometrische Datierung des Unter-Eozäns von Helmstedt (SE-Niedersachsen). Zeitschrift der Deutschen Geologischen Gesellschaft 1995; 146: 450–457.

39. Köthe A. Dinozysten-Zonierung im Tertiär Norddeutschlands. Revue de Paléobiologie 2003; 22: 895–923.

40. Pflug HD. Palynologie und Stratigraphie der eozänen Braunkohlen von Helmstedt. Paläontologische Zeitschrift 1952; 26: 112–137.

41. Pflug HD. Palyno-Stratigraphie des Eozän/Oligozän im Raum von Helmstedt, in Nordhessen und im südlichen Anschlussbereich. In: Tobien H, editor. Nordwestdeutschland im Tertiär. Beiträge zur Regionalen Geologie der Erde 18: Berlin, Stuttgart: Gebrüder Borntraeger; 1986. pp. 567–582.

42. Haq BU, Hardenbol J, Vail PR. Mesozoic and Cenozoic chronostratigraphy and cycles of sea-level change. In: Wilgus CK, Hastings BS, Kendall CGSG, Posamentier HW, Ross CA, Van Wagoner JC, editors. Sea-level changes: an integrated approach. Tulsa, USA: SEPM (Society for Sedimentary Geology) Special Publication 42; 1988. pp. 71–108.

43. Laskar J, Fienga A, Gastineau M, Manche H. La2010. A new orbital solution for the long term motion of the Earth. Astronomy & Astrophysics 2011; 532, A89: 1–15.

44. Riegel W, Wilde V. An early Eocene Sphagnum bog at Schöningen, northern Germany. International Journal of Coal Geology 2016; 159: 57–70.

45. Bujak JP, Brinkhuis H. Global warming and dinocyst changes across the Paleocene/Eocene Epoch boundary. In: Aubry MP, Lucas SG, Berggren W, editors. Late Paleocene–early Eocene climatic and biotic events in the marine and terrestrial records. New York, USA: Columbia University Press; 1998. pp. 277–295.

46. Crouch EM, Heilmann-Clausen C, Brinkhuis H, Morgans HE, Rogers KM, Egger H, et al. Global dinoflagellate event associated with the late Paleocene thermal maximum. Geology 2001; 29: 315–318.

47. Heilmann-Clausen C, Nielsen OB, Gersner F. Lithostratigraphy and depositional environments in the Upper Paleocene and Eocene of Denmark, Bulletin of the Geological Society of Denmark. 1985; 33: 287–323.

48. Iakovleva AI, Brinkhuis H, Cavagnetto C. Late Palaeocene–Early Eocene dinoflagellate cysts from the Turgay Strait, Kazakhstan; correlations across ancient seaways, Palaeogeography, Palaeoclimatology, Palaeoecology 2001; 172: 243–268.

49. Sluijs A, Brinkhuis H. A dynamic climate and ecosystem state during the Paleocene-Eocene Thermal Maximum: inferences from dinoflagellate cyst assemblages on the New Jersey Shelf. Biogeosciences 2009; 6: 1755–1781.

50. Sluijs A, Schouten S, Pagani M, Woltering M, Brinkhuis H, Damsté JSS, et al. Subtropical Arctic Ocean temperatures during the Palaeocene/Eocene thermal maximum. Nature 2006; 441: 610–613.

51. Sluijs A, Brinkhuis H, Schouten S, Bohaty SM, John CM, Zachos JC, et al. Environmental precursors to rapid light carbon injection at the Palaeocene/Eocene boundary. Nature 2007; 450: 1218–1221.

52. Williams GL, Damassa SP, Fensome RA, Guerstein GR. Wetzeliella and its allies — The ‘Hole’ Story: A Taxonomic Revision of the Paleogene Dinoflagellate Subfamily Wetzelielloideae. Palynology 2015; 39: 289–344.

53. Heilmann-Clausen C. Observations of the dinoflagellate *Wetzeliella* in Sparnacian facies (Eocene) near Epernay, France, and a note on tricky acmes of *Apectodinium*. Proceedings of the Geologists’ Association. 2018. https://doi.org/10.1016/j.pgeola.2018.06.001

54. Kaiser ML, Ashraf R. Gewinnung und Präparation fossiler Pollen und Sporen sowie anderer Palynomorphae unter besonderer Berücksichtigung der Siebmethode. Geologisches Jahrbuch 1974; 25, 85–114.

55. Thomson PW, Pflug H. Pollen und Sporen des mitteleuropäischen Tertiärs. Gesamtübersicht über die stratigraphisch und paläontologisch wichtigen Formen. Palaeontographica B 1953; 94: 1–138.

56. Krutzsch W, Vanhoorne R. Die Pollenflora von Epinois und Loksbergen in Belgien. Palaeontographica B 1977; 163: 1–110.

57. Thiele-Pfeiffer H. Die Mikroflora aus dem mitteleozänen Ölschiefer von Messel bei Darmstadt. Palaeontographica B 1988; 211: 1–86.

58. Nickel B. Die mitteleozäne Mikroflora von Eckfeld bei Manderscheid/Eifel. Mainzer Naturwissenschaftliches Archiv Beiheft 1996; 18: 1–121.

59. ter Braak CJF, Looman CWN. Regression. In: Jongman RHG, Ter Braak CJF, Tongeren OFR, editors. Data Analysis in Community and Landscape Ecology. Cambridge, UK: Cambridge University Press; 1995. pp. 29–77.

60. Juggins S. C2 Software for ecological and palaeoecological data analysis and visualization. User Guide Version, 1.5. Newcastle upon Tyne: Newcastle University; 2007.

61. Lenz OK, Wilde V. Changes in Eocene plant diversity and composition of vegetation: the lacustrine archive of Messel (Germany). Paleobiology 2018; 44: 709–735.

62. Moshayedi M, Lenz OK, Wilde V, Hinderer M. The recolonization of volcanically disturbed Eocene habitats of Central Europe: The maar lakes of Messel and Offenthal (SW Germany) compared. Palaeobiodiversity and Palaeoenvironments 2020. doi: 10.1007/s12549-020-00425-4.

63. Bray JR, Curtis JT. An ordination of the upland forest communities of southern Wisconsin. Ecological Monographs 1957; 27: 325–349.

64. Cottam G, Goff FG, Whittaker RH. Wisconsin Comparative Ordination. In: Whittaker RH, editor. Ordination of Plant Communities. Handbook of Vegetation Science 5-2. Dordrecht: Springer; 1978. pp. 185–213.

65. Gauch HG, Scruggs WM. Variants of polar ordination. Vegetatio 1979; 40: 147–153.

66. Oksanen J. Multivariate Analysis of Ecological Communities in R. 2005 [cited 2020 September 10]. Available from: http://ubio.bioinfo.cnio.es/Cursos/CEUMDA07practicals/Furtherreading/Oksanen2005/

67. Mander L, Kürschner WM, McElwain JC. An explanation for conflicting records of Triassic–Jurassic plant diversity. Proceedings of the National Academy of Sciences of the United States of America 2010; 107: 15351–15356.

68. Hammer Ø, Harper DAT, Ryan PD. PAST: paleontological statistics software package for education and data analysis. Palaeontologia Electronica 4; 2001. [cited 2020 September 10]. Available from: http://www.palaeo-electronica.org/2001_1/past/issue1_01.htm.

69. 69. Hair JF, Black WC, Babin BJ, Anderson RE. Multivariate Data Analysis. Seventh Edition. Prentice Hall, Upper Saddle River, New Jersey; 2010.

70. Minchin PR. An evaluation of the relative robustness of techniques for ecological ordination. Vegetatio 1987; 69: 89–107.

71. Jardine PE, Harrington GJ. The Red Hills Mine palynoflora: A diverse swamp assemblage from the Late Paleocene of Mississippi, USA. Palynology 2008; 32: 183–204.

72. Ghilardi B, O’Connell M. Fine resolution pollen analytical study of Holocene woodland dynamics and land use in north Sligo, Ireland. Boreas 2013; 42: 623–649.

73. Broothaerts N, Verstraeten G, Kasse C, Bohncke S, Notebaert B, Vandenberghe J. Reconstruction and semi-quantification of human impact in the Dijle catchment, central Belgium: a palynological and statistical approach. Quaternary Science Reviews 2014; 102: 96–110.

74. Borsch T, Wilde V. Pollen variability within species, populations, and individuals, with particular reference to Nelumbo nucifera. In: Harley M, Blackmore S, Morton C, editors. Pollen and Spores: Morphology and Biology. London: Royal Botanic Gardens, Kew; 2000. pp. 285–299.

75. Colwell RK, Chao A, Gotelli NJ, Lin SY, Mao CX, Chazdon RL, et al. Models and estimators linking individual-based and sample-based rarefaction, extrapolation, and comparison of assemblages. Journal of Plant Ecology 2012; 5: 3–21.

76. Colwell RK. EstimatesS: Statistical estimation of species richness and shared species from samples. Version 9. 2013. Available from: http://viceroy.eeb.uconn.edu/EstimateS/

77. Keen HF, Gosling WD, Hanke F, Miller CS, Montoya E, Valencia BG, et al. A statistical sub-sampling tool for extracting vegetation community and diversity information from pollen assemblage data. Palaeogeography, Palaeoclimatology, Palaeoecology 2014; 408: 48–59.

78. Whittaker RH. Evolution and measurement of species diversity. Taxon 1972; 21: 213– 251.

79. Harrington GJ, Jaramillo CA. Paratropical floral extinction in the Late Palaeocene–Early Eocene. Journal of the Geological Society London 2007; 164: 323–332.

80. Birks HJB, Felde VA, Bjune AE, Grytnes JA, Seppä H, Giesecke T. Does pollen-assemblage richness reflect floristic richness? A review of recent developments and future challenges. Review of Palaeobotany and Palynology 2016; 228: 1–25.

81. Ellison AM. Partitioning diversity. Ecology 2010; 91: 1962–1963.

82. Beck J, Holloway JD, Schwanghart W. Undersampling and the measurement of beta diversity. Methods in Ecology and Evolution 2013; 4: 370–382.

83. Whittaker RH. Vegetation of the Siskiyou mountains, Oregon and California. Ecological Monographs 1960; 30: 279–338.

84. Koleff P, Gaston KJ, Lennon JJ. Measuring beta diversity for presence–absence data. Journal of Animal Ecology 2003; 72: 367–382.

85. Gotelli NJ, Colwell RK. Estimating species richness. In Magurran AE, McGill BJ, editors. Biological diversity. Frontiers in measurement and assessment. Oxford University Press, New York; 2010. pp. 39–54.

86. Smith B, Wilson JB. A consumer’s guide to evenness indices. Oikos 1996; 76: 70–82.

87. Vogt W. Makropetrographischer Flözaufbau der rheinischen Braunkohle und Brikettiereigenschaften der Lithotypen. Fortschritte in der Geologie von Rheinland und Westfalen 1981; 29: 73–93.

88. Faegri K, Iversen J, Kaland PE, Krzywinsi, K. Textbook of pollen analysis. 4th ed. Chichester. New York, Brisbane, Tokyo, Singapore: John Wiley & Sons; 1989.

89. Inglis GN, Collinson ME, Riegel W, Wilde V, Robson BE, Lenz OK, et al. Ecological and biogeochemical change in an early Paleogene peat-forming environment: Linking biomarkers and palynology, Palaeogeography, Palaeoclimatology, Palaeoecology 2015; 438: 245–255.

90. Gastaldo RA, Allen GP, Huc AY. Detrital peat formation in the tropical Mahakam River delta, Kalimantan, eastern Borneo: Sedimentation, plant composition, and geochemistry. In: Cobb JC, Blaine C, editors. Modern and Ancient Coal-Forming Environments Mires. USA: Geological Society of America Special Paper 286; 1993. pp. 107–118.

91. Gastaldo, RA. The genesis and sedimentation of phytoclasts with examples from coastal environments. In: Traverse A, editor. Sedimentation of Organic Particles. Cambridge, UK: Cambridge University Press; 1994. pp. 103–127.

92. Krebs CJ. Ecological methodology. New York, NY: Harper and Row Publishers Inc.; 1989.

93. Cohen AD. Peats from the Okefenokee Swamp-marsh complex. Geoscience and Man 1975; 11: 123–131.

94. Willard DA, Weimer LM, Riegel W. Pollen assemblages as paleoenvironmental proxies in the Florida Everglades. Review of Palaeobotany and Palynology 2001; 113: 213–235.

95. Schneck T. Umweltrekonstruktion mit Hilfe der Palynologie: eine Studie über bisher ungenutzte Potentiale. PhD Thesis, Eberhard Karls Universität Tübingen. 2006. Available from: http://hdl.handle.net/10900/48942

96. Smith AHV. The sequence of microspore assemblages associated with the occurrence of crassidurite in coal seams of Yorkshire. Geological Magazine 1957; 94: 345–363.

97. Smith AHV. The palaeoecology of carboniferous peats based on the miospores and petrography of bitmminous coals. Proceedings of the Yorkshire Geological Society 1962; 33: 423–474.

98. Smith AVH. Seam profiles and seam characters. In: Murchison D, Westoll TS, editors. Coal and coal-bearing strata. New York: Elsevier; 1968. pp. 31–40.

99. Dai S, Bechtel A, Eble CF, Flores RM, French D, Graham IT, et al. Recognition of peat depositional environments in coal: A review. International Journal of Coal Geology 2020. doi: 10.1016/j.coal.2019.103383

100. Daubenmire RF. Plant Geography with special reference to North America. New York, NY: Academic Press; 1978.

101. Box EO. Warm-Temperate Deciduous Forests of Eastern North America. In: Box E, Fujiwara K, editors. Warm-Temperate Deciduous Forests around the Northern Hemisphere. Geobotany Studies (Basics, Methods and Case Studies). Cham: Springer; 2015. pp. 225–255.

102. Teichmüller M. Rekonstruktion verschiedener Moortypen des Hauptflözes der niederrheinischen Braunkohle. Fortschritte in der Geologie des Rheinlandes und Westfalen 1958; 2: 539 612.

103. Teichmüller M. The genesis of coal from the viewpoint of coal petrology. International Journal of Coal Geology 1989; 12: 1 87.

104. Ivanov DA, Koleva-Rekalova R. Palynological and sedimentological data about late Sarmatian palaeoclimatic changes in the Fore-Carpathian and Euxinian Basins (northern Bulgaria). Acta Palaeobotanica Supplementum 1999; 2: 307–313.

105. Karon R. Palinofacies in the Turów open pit (SW Poland). Acta Palaeobotanica Supplementum 1999; 2: 315–317.

106. Riegel W, Bode T, Hammer J, Hammer-Schiemann G, Lenz, O, Wilde V. The Palaeoecology of the lower and middle Eocene at Helmstedt, northern Germany: a study in contrasts: Acta Palaeobotanica Supplementum 1999; 2: 349–358.

107. Holdgate G, Wallace M, O’Connor M, Korasidis V, Lieven U. The origin of lithotype cycles in Oligo-Miocene brown coals from Australia and Germany. International Journal of Coal Geology 2016; 166: 47–61.

108. Lenz OK, Riegel W. Isopollen maps as a tool for the reconstruction of a coastalswamp from the Middle Eocene at Helmstedt (Northern Germany). Facies 2001; 45: 177–194.

109. Wilde V, Lenz OK, Riegel W. Mangrove structure and development in the Lower and Middle Eocene of Helmstedt, northern Germany. Terra Nostra 2008; 2: 306–307.

110. Natlandsmyr B, Hjelle KL. Long-term vegetation dynamics and land-use history: Providing a baseline for conservation strategies in protected *Alnus glutinosa* swamp woodlands. Forest Ecology and Management 2016; 372: 78 92.

111. Houston DT, de Rigo D, Caudullo G. Alnus incana in Europe: distribution, habitat, usage and threats. In: San-Miguel-Ayanz J, de Rigo D, Caudullo G, Houston DT, Mauri A, editors. European Atlas of Forest Tree Species. Luxembourg: Publication Office of the European Union; 2016: pp. e01ff87+.

112. Fralish JS, Franklin SB. Taxonomy and Ecology of Woody Plants in North American Forests (Excluding Mexico and Subtropical Florida). New York, NY: John Wiley & Sons; 2002.

113. Spackman W, Riegel WL, Dolsen CP. Geological and biological interactions in the swamp-marsh complex of Southern Florida. In: Dapples EC, Hopkins ME, editors. Environments of Coal Deposition. USA: Geological Society of America Special Paper 114; 1969. pp. 1–35.

114. Cohen AD. Possible influences of subpeat topography and sediment type upon the development of the Okefenokee swamp-marsh complex of Georgia. Southeastern Geology 1974; 15: 141–151.

115. Hamilton DB. Plant succession and the influence of disturbance in Okefenokee Swamp. In: Cohen AD, Casagrande DJ, Andrejko MJ, Best GR, editors. The Okefenokee Swamp: Its Natural History, Geology, and Geochemistry. Los Alamos: Wetland Surveys; 1984. pp. 86– 106.

116. Ewel KC. Swamps. In: Myers RL, Ewel JJ, editors. Ecosystems of Florida. Orlando: University of Central Florida Press; 1990. pp. 281–323.

117. Gastaldo RA, Staub JR. A mechanism to explain the preservation of leaf litter lenses in coals derived from raised mires. Palaeogeography, Palaeoclimatology, Palaeoecology 1999; 149: 1–14.

118. Punt W, Marks A, Hoen PP. The Northwest European Pollen Flora, 66. Myricaceae. Review of Palaeobotany and Palynology 2002; 123: 99–105

119. Grimsson F., Grimm GW, Meller B, Bouchal JM, Zetter R. Combined LM and SEM study of the middle Miocene (Sarmatian) palynoflora from the Lavanttal Basin, Austria: part IV.Magnoliophyta 2 – Fagales to Rosales. Grana 2016; 55: 101–163.

120. Simpson MJA., Macintosh DF, Cloughley JB., Stuart AE. Past, present and future utilisation of *Myrica gale* (Myricaceae). Economic Botany 1996; 50: 122–129.

121. Skene KR, Sprent JI, Raven JA, Herdman L. Myrica gale L. Biological flora of the British Isles. Journal of Ecology 2000; 88: 1079–1094.

122. van Breemen N. How *Sphagnum* bogs down other plants. Trends in Ecology & Evolution 1995; 10: 270–275.

123. Clymo R. 1984. The limits to peat bog growth. Philosophical Transactions of the Royal Society B 1984; 303: 605–654.

124. Page SE, Rieley JO, Shotyk W, Weiss D. Interdependence of peat and vegetation in tropical swamp forest. Philosophical Transactions of the Royal Society B 1999; 354: 1885– 1897.

125. Cohen AD, Andrejko MJ, Spackman W, Corvinus D. Peat deposits of the Okefenokee Swamp. In: Cohen AD, Casagrande DJ, Andrejko MJ, Best GR. editors. The Okefenokee Swamp: Its Natural History, Geology, and Geochemistry. Los Alamos: Wetland Surveys; 1984. pp. 493–553.

126. Moore PD. Ecological and hydrological aspects of peat formation. In: Scott AC, editor. Coal and Coal-bearing Strata: Recent Advances. London, UK; Geological Society London Special Publication 32; 1987. pp. 7–16.

127. Korasidis VA, Wallace MW, Tosolini AMP, Hill RS. The origin of floral lagerstätten in coals. Palaios 2020; 35: 22–-36.

128. Daly RJ, Jolley DW. What was the nature and role of Normapolles angiosperms? A case study from the earliest Cenozoic of Eastern Europe. Palaeogeography, Palaeoclimatology, Palaeoecology 2015; 418: 141–149.

129. Loftin SS, Guyette MQ, Wetzel PR. Evaluation of Vegetation-Fire Dynamics in the Okefenokee National Wildlife Refuge, Georgia, USA, with Bayesian Belief Networks. Wetlands 2018; 38: 819 834.

130. Cypert E. The effect of fires in the Okefenokee Swamp in 1954 and 1955. The American Midland Naturalis. 1961; 66: 485–503.

131. Cohen AD. Petrography and paleoecology of Holocene peats from the Okefenokee swamp-marsh complex of Georgia. Journal of Sedimentary Petrology 1974; 44: 716–726.

132. Izlar, RL. Some comments on fire and climate in the Okefenokee swamp-marsh complex. In: Cohen AD, Casagrande DJ, Andrejko MJ, Best GR, editors. The Okefenokee swamp: its natural history, geology and geochemistry. Los Alamos: Wetland Surveys; 1984. pp. 70–85.

133. Hofmann CC, Zetter R. Palynological investigations of the Krappfeld area, Palaeocene/Eocene, Carinthia (Austria). Palaeontographica B 2001; 259: 47–64

134. Zetter R., Hofmann CC. New aspects on the palyno-flora of the lowermost Eocene (Krappfeld, Carinthia). In: Piller WE, Rasser MW, editors. Paleogene of the Eastern Alps. Wien: Verlag der Österreichischen Akademie der Wissenschaften 12; 2001. pp. 473–507.

135. Harrington GJ. Impact of Paleocene/Eocene Greenhouse Warming on North American Paratropical Forests. Palaios 2001; 16: 266–278.

136. Harrington GJ. Geographic patterns in the floral response to Paleocene–Eocene warming. In: Wing SL, Gingerich PD, Schmitz B, Thomas E, editors. Causes and consequences of globally warm climates in the early Paleogene. USA: Geological Society of America Special Paper 369; 2003. pp. 381–393.

137. Page SE, Rieley JO, Wust R. Chapter 7. Lowland tropical peatlands of Southeast Asia. In: Martini IP, Martinez Cortizas A, Chesworth W, editors. Peatlands-Evolution and Records of Environmental and Climate Changes. Elsevier: Developments in Earth Surface Processes 9; 2006. pp. 145–172.

138. Phillips S, Rouse GE, Bustin RM. Vegetation zones and diagnostic pollen profiles of a coastal peat swamp, Bocas del Toro, Panamá. Palaeogeography, Palaeoclimatology, Palaeoecology 2001; 128: 301–338.

139. Inglis GN, Collinson ME, Riegel W, Wilde V, Farnsworth A, Lunt DJ, et al. Mid-latitude continental temperatures through the early Eocene in western Europe. Earth and Planetary Science Letters 2017; 460: 86–96.

140. Greenwood DR, Basinger JF. The paleoecology of high-latitude Eocene swamp forests from Axel Heiberg Island, Canadian High Arctic. Review of Palaeobotany and Palynology 2013; 81: 83–97.

141. Eldrett JS, Greenwood DR, Polling M, Brinkhuis H, Sluijs A. A seasonality trigger for carbon injection at the Paleocene–Eocene Thermal Maximum. Climate of the Past 2014; 10: 759–769.

142. Weijers JWH, Schouten S, Sluijs A, Brinkhuis H, Sinninghe Damsté JS. Warm arctic continents during the Palaeocene–Eocene thermal maximum. Earth and Planetary Science Letters 2007; 261: 230–238.

143. Wing SL, Bao H, Koch PL. An early Eocene cool period? Evidence for continental cooling during the warmest part of the Cenozoic. In: Huber BT, MacLeod KG, Wing SL, editors. Warm Climates in Earth History. Cambridge, UK: Cambridge University Press; 2000. pp. 197–237.

144. Campbell EO. The restiad peat bogs at Motumaoho and Moanatuatua. Transactions of the Royal Society of New Zealand 1964; 2: 219–227.

145. Campbell EO. Peat deposits of northern New Zealand as based on identification of plant fragments in the peat. Proceedings of the New Zealand Ecological Society 1975; 22: 57–60.

146. Campbell EO. Mires of Australasia. In: Gore AJP, editor. Mires: swamp, bog, fen and moor ecosystems of the world 4A. Amsterdam, Oxford, New York: Elsevier Scientific Publishing Company; 1983. pp. 153–180.

147. Clarkson BR, Schipper LA, Lehmann A. Vegetation and peat characteristics in the development of lowland restiad peat bogs, North Island, New Zealand. Wetlands 2004; 24: 133–151.

148. Benson D, Baird IRC. Vegetation, fauna and groundwater interrelations in low nutrient temperate montane peat swamps in the upper Blue Mountains, New South Wales. Cunninghamia 2012; 12: 267–307

149. Fairfax R, Lindsay R. An overview of the patterned fens of Great Sandy Region, far eastern Australia. Mires and Peat 2019; 24: 1–18. doi: 10.19189/MaP.2018.OMB.369.

150. Nichols DJ, Pocknall DT. Relationships of palynofacies to coal-depositional environments in the upper Paleocene of the Gulf Coast Basin, Texas, and the Powder River Basin, Montana and Wyoming. Traverse A, editor. Sedimentation of Organic Particles. Cambridge University Press, Cambridge; 1994. pp. 217–237.

151. Innes JB, Simmons I. Mid Holocene charcoal stratigraphy, fire history and palaeoecology at North Gill, North York Moors, UK. Palaeogeography, Palaeoclimatology, Palaeoecology 2000; 164: 151–165.

152. Nichols GJ, Cripps JA, Collinson ME, Scott AC. Experiments in waterlogging and sedimentology of charcoal: results and implications. Palaeogeography, Palaeoclimatology, Palaeoecology 2000; 164: 43–56.

153. Scott AC, Glasspool IJ. Observations and experiments on the origin and formation of inertinite group macerals. International Journal of Coal Geology 2007; 70: 53–66.

154. Scott AC. Charcoal recognition, taphonomy and uses in palaeoenvironmental analysis. Palaeogeography, Palaeoclimatology, Palaeoecology 2010; 291: 11–39.

155. Clark JS. Particle motion and the theory of charcoal analysis: source area, transport, deposition, and sampling. Quaternary Research 1988; 30: 67–80.

156. Clark JS, Lynch J, Stocks BJ, Goldammer, JG. Relationships between charcoal particles in air and sediments in west-central Siberia. The Holocene 1998; 8: 19–29.

157. Collinson ME, Hooker JJ, Gröcke DR. Cobham Lignite Bed and penecontemporaneous macrofloras of southern England: A record of vegetation and fire across the Paleocene-Eocene Thermal Maximum. Geological Society of America, Special Paper 2003; 369: 333 – 349.

158. Kurtz AC, Kump LR, Arthur MA, Zachos JC, Paytan A. Early Cenozoic decoupling of the global carbon and sulphur cycles. Palaeoceanography 2003; 18: 1090–1104.

159. Moore EA, Kurtz AC. Black carbon in Paleocene–Eocene boundary sediments: A test of biomass combustion as the PETM trigger. Palaeogeography, Palaeoclimatology, Palaeoecology 2008; 267: 147–152.

160. Korasidis VA, Wallace MW, Wagstaff BE, Hill RS. Evidence of fire adaption in Australian Cenozoic rainforests. Palaeogeography, Palaeoclimatology, Palaeoecology 2019; 516: 35–43.

