## Supplementary material for "Greenhouse conditions in lower Eocene coastal wetlands? – Lessons from Schöningen, Northern Germany": S1_Table

**S1 Table**: **Complete list of palynomorphs from the studied sections N, S1 and S2 of Seam 1 of the Schöningen Formation including their systematic affinities.** In the left column the 45 “variables” are presented, which were used for the pollen diagrams and statistical analyses (cluster analysis, non-metric multidimensional scaling). The list includes 190 different spore and pollen species, which were used for analyses of palynological morpho-diversity.

| Variable (used in pollen diagrams and for statistical analyses) | Palynomorphs | Family/Genus |
| --- | --- | --- |
| **Spores** | | |
| *Sphagnum*-type spores | *Distancorisporis* sp*.* Krutzsch in Döring et al. 1966) Srivastava, 1972 | Sphagnaceae, *Sphagnum*? |
|  | *Tripunctisporis* spp. (Krutzsch in Döring et al. 1966) Herngreen et al. 1986, 2 morpho-types | Sphagnaceae, *Sphagnum*? |
|  | *Sphagnumsporites* sp*.* (Raatz 1937) | Sphagnaceae, *Sphagnum* |
| *Leiotriletes* group | *Leiotriletes adriennis* (R. Potonié & Gelletich 1933) Krutzsch 1959 | Schizaeaceae, *Lygodium*? |
|  | *Leiotriletes microadriennis* Krutzsch 1959 | Schizaeaceae, *Lygodium* |
|  | *Leiotriletes paramaximus* Krutzsch 1959 | Schizaeaceae? |
|  | *Leiotriletes dorogensis* Kedves 1960 | Schizaeaceae? |
|  | *Leiotriletes triangulus* (Mürriger & Pflug 1952) Krutzsch 1962 | ? |
|  | *Intrapunctisporis gracilioides* Krutzsch & Vanhoorne 1977 | ? |
|  | *Toroisporis (Toroisporis) megatorus* Krutzsch & Vanhoorne 1977 | ? |
|  | *Toroisporis (Toroisporis) longitorus* Krutzsch 1959 | ? |
|  | *Toroisporis (Toroisporis) eocaenicus* Kedves 1966 | ? |
|  | *Toripunctisporis* sp*.* | ? |
|  | *Concavisporites pseudopartitus* Krutzsch 1959 | ? |
| other trilete spores | *Neogenisporis neogenicus* Krutzsch 1962 | Gleicheniaceae, Cyatheaceae |
|  | *Neogenisporis pseudoneddeni (*Krutzsch 1959) Krutzsch 1962 | Gleicheniaceae, Cyatheaceae |
|  | *Neogenisporis sp.* | Gleicheniaceae, Cyatheaceae |
|  | *Cicatricosisporites dorogensis* R. Potonié & Gelletich 1933 | Schizaeaceae, *Mohria, Anemia* |
|  | *Foveotriletes crassifovearis* Krutzsch 1962 | Ophioglossaceae, Botrychium |
|  | *Corrugatisporites corruvallatus (*Krutzsch 1967) Nagy 1985 | Ophioglossaceae, Botrychium |
|  | *Trilites multivallatus* (Pflug 1953) Krutzsch 1959 | Schizaeaceae?, Lygodiaceae? |
|  | *Trilites* sp. | ? |
|  | *Goczanisporis baculatus* Krutzsch 1967 | ? |
|  | *Retitriletes rueterbergensis* Krutzsch 1963 | Lycopodiaceae, *Lycopodium* |
|  | *Camarozonosporites heskemensis* (Pflanzl 1955) Krutzsch 1959, 2 morpho-types | Lycopodiaceae, *Lycopodium* |
|  | *Baculatisporis primarius* (Wolff 1934) Thomson & Pflug 1953 | Osmundaceae, *Osmunda* |
|  | *Tegumentisporis tegumentis (*Krutzsch 1959) Krutzsch 1963 | Selaginellaceae, *Selaginella* |
| *Laevigatosporites* spp. | *Laevigatosporites haardtii* (R. Potonié & Venitz 1934) Thomson & Pflug 1953 | Polypodiaceae |
|  | *Laevigatosporites discordatus* Pflug 1953 | Polypodiaceae |
|  | *Laevigatosporites pseudodiscordatus* Krutzsch 1959 | Polypodiaceae |
| *Verrucatosporites* spp. | *Verrucatosporites favus* (R. Potonié 1931) Thomson & Pflug 1953 | Polypodiaceae, *Polypodium* |
|  | *Verrucatosporites microfavus* Thiele-Pfeiffer 1988 | Polypodiaceae |
| other monolete spores | *Punctatosporites palaeogenicus* Krutzsch 1959 | Polypodiaceae |
| **Pollen** | | |
| Bisaccates | *Pityosporites labdacus* (R. Potonié 1931) Thomson & Pflug 1953 *(Pinus-type)* | Pinaceae, *Pinus* |
|  | *Pityosporites labdacus* (R. Potonié 1931) Thomson & Pflug 1953 *(Cathaya-type)* | Pinaceae, *Cathaya* |
| Inaperturate pollen | *Inaperturopollenites dubius* (R. Potonié & Venitz 1934) Thomson & Pflug 1953 | Cupressaceae |
|  | *Inaperturopollenites hiatus* (R. Potonié 1931) Thomson & Pflug 1953 | Cupressaceae |
|  | *Inaperturopollenites concedipites* (Wodehoise 1933) Krutzsch 1971 | Cupressaceae, *Glyptostrobus*? |
|  | *Inaperturopollenites magnus* (R. Potonié 1934) Thomson & Pflug 1953 | Cupressaceae?, Pinaceae? |
|  | *Inaperturopollenites* sp*.* | Cupressaceae |
|  | *Cupressacites bockwitzensis* Krutzsch 1971 | Cupressaceae |
|  | *Sciadopityspollenites* sp*.* | Cupressaceae, *Sciadopitys*? |
|  | *Sequoiapollenites* spp*.,* 2 morpho-types | Cupressaceae, *Sequoia* |
| *Monocolpopollenites tranquillus* | *Monocolpopollenites tranquillus* (R. Potonié 1934) Thomson & Pflug 1953 | Arecaceae, *Phoenix* |
| *Dicolpopollis kockeli* | *Dicolpopollis kockeli* Pflanzl 1956 | Arecaceae, *Calamus* |
| *Arecipites* spp. | *Arecipites convexus* (Thiergart 1934) Krutzsch 1970 | Arecaceae, *Sabal*?, *Pseudophoenix*?*, Areca*? |
|  | *Arecipites longicolpatus* Krutzsch 1970 | Arecaceae, *Sabal*?*, Pseudophoenix*?*, Areca*? |
| other monocolpate pollen | *Liliacidites monosulcoides (*Krutzsch 1970) Kohlman-Adamska & Ziembinska-Tworzydlo 2014 | Asparagaceae?, Amaryllidaceae?, Araceae?, Iridaceae?, Liliaceae? |
|  | *Liliacidites konzalovae* Kohlman-Adamska & Ziembinska-Tworzydlo 2014 | Asparagaceae?, Amaryllidaceae?, Araceae?, Iridaceae?, Liliaceae? |
| *Milfordia* spp*.* | *Milfordia hungarica* (Kedves 1965) Krutzsch & Vanhoorne 19977 in Krutzsch 1970 | Restionaceae, *Lyginia, Restio* |
|  | *Milfordia incerta* (Thomson & Pflug 1953) Krutzsch 1961 | Restionaceae, *Hypolaena* |
| *Sparganiaceaepollenites* spp. | *Sparganiaceaepollenites cuvillieri* (Cavagnetto 1966) Roche 1968 | Sparganiaceae, Typhaceae |
|  | *Sparganiaceaepollenites sparganioides* (Meyer 1956) Krutzsch 1970 | Sparganiaceae, Typhaceae |
| *Emmapollis pseudoemmaensis* | *Emmapollis pseudoemmaensis* Krutzsch 1970 | Chloranthaceae, *Ascarinopsis*?*, Ascarina*? |
| Normapolles elements | *Basopollis orthobasalis* (Pflug 1953a) Pflug 1953b | ? |
|  | *Basopollis atumescens* (Thomson & Pflug 1953) Pflug 1953 | ? |
|  | *Nudopollis terminalis hastaformis* (Krutzsch 1954) Krutzsch in Góczán, Groot, Krutzsch & Pacltová 1967 | ? |
|  | *Trudopollis* sp*.* | ? |
| *Thomsonipollis magnificus* | *Thomsonipollis magnificus* (Thomson & Pflug 1953) Krutzsch 1960 | ? |
|  | *Thomsonipollis magnificoides* Krutzsch 1960 | ? |
| *Pistillipollenites mcgregorii* | *Pistillipollenites mcgregorii* Rouse 1962 | Gentianaceae |
| *Plicapollis pseudoexcelsus* | *Plicapollis pseudoexcelsus* (Krutzsch 1957) Krutzsch 1961 *turgidus* Pflug 1953 | Juglandales |
|  | *Plicapollis pseudoexcelsus* (Krutzsch 1957) Krutzsch 1961 *semiturgidus* Pflug 1953 | Juglandales |
|  | *Plicapollis pseudoexcelsus* (Krutzsch 1957) Krutzsch 1961 *microturgidus* Pflug 1953 | Juglandales |
| *Pompeckjoidaepollenites subhercynicus* | *Pompeckjoidaepollenites subhercynicus* (Krutzsch 1954) Krutzsch in Góczán*,* Groot, Krutzsch & Pacltová 1967, 2 morpho-types | ? (Normapolles) |
| *Interpollis* sp. | *Interpollis supplingensis* (Pflug 1953) Krutzsch 1961 | ? (Normapolles) |
| *Platycaryapollenites* spp*.* | *Platyacaryapollenites saxonis* Krutzsch 1969 | Juglandaceae, *Platycarya* |
|  | *Platycaryapollenites miocaenicus* Nagy 1969 | Juglandaceae, *Platycarya* |
|  | *Platycaryapollenites levis* (R. Potonié 1931) Krutzsch 1969 | Juglandaceae, *Platycarya* |
|  | *Platycaryapollenites semicyclus* (Krutzsch & Vanhoorne 1977) Thiele-Pfeiffer 1988 | Juglandaceae, *Platycarya*? |
|  | *Platycaryapollenites platycaryoides* (Roche 1969) Kedves 1982 | Juglandaceae, *Platycarya* |
| *Plicatopollis plicatus* group | *Plicatopollis plicatus* (R. Potonié 1934) Krutzsch 1962 | Juglandaceae |
|  | *Plicatopollis hungaricus* Kedves 1974 | Juglandaceae |
|  | *Plicatopollis lunatus* Kedves 1974 | Juglandaceae |
|  | *Plicatopollis* sp*.* | Juglandaceae |
|  | *Pseudoplicapollis palaeocaenicus* Krutzsch in Góczán*,* Groot, Krutzsch & Pacltová 1967 | Juglandaceae? |
| *Triporollenites robustus/rhenanus* group | *Triporopollenites robustus* (Mürriger & Pflug 1951) Thomson & Pflug 1953 | Myricaceae |
|  | *Triporopollenites undulates* Pflug 1953 | Myricaceae |
|  | *Triporopollenites crassus* Krutzsch & Vanhoorne 1977 | Myricaceae |
|  | *Triporopollenites rhenanus* (Thoms. in R. Pot., Thoms. & Thierg. 1950) Th. & Pf. 1953 | Myricaceae, Betulaceae?, *Ostrya*? |
|  | *Triporopollenites intrastructurus* Krutzsch & Vanhoorne 1977 | Betulaceae |
| *Triatriopollenites bituitus/rurensis* group | *Triatriopollenites excelsus* (R. Potonié 1931) Thomson & Pflug 1953 | Myricaceae, Juglandaceae? |
|  | *Triatriopollenites excelsus* (R. Potonié 1931) Thomson & Pflug 1953 *minor* Pflug 1953 | Myricaceae, Juglandaceae? |
|  | *Triatriopollenites rurensis* Thomson & Pflug 1953 | Myricaceae, *Myrica* |
|  | *Triatriopollenites bituitus* (R. Potonié 1931) Thomson & Pflug 1953 | Myricaceae, *Myrica* |
|  | *Triatriopollenites sibiricus* (Gladkova 1965) Kedves 1974 | Myricaceae |
|  | *Triatriopollenites pseudoroboratus* Krutzsch & Vanhoorne 1977 | Myricaceae? |
| *Momipites punctatus* | *Momipites punctatus* (R. Potonié 1931) Nagy 1969 | Juglandaceae, *Engelhardia*?, *Oreomunnea*?, *Alfaroa*? |
| *Pentapollis pentangulus* | *Pentapollis pentangulus* (Pflug 1953) Krutzsch 1957 | ? |
| *Labrapollis labraferus* | *Labrapollis labraferus* (R. Potonié 1931) Krutzsch 1968 | ? |
|  | *Labrapollis rotundoides* Krutzsch & Vanhoorne 1977 | ? |
| *Intratriporopollenites* spp. | *Intratriporopollenites giganteus* Krutzsch & Vanhoorne 1977 | Tiliaceae |
|  | *Intratriporopollenites insculptus* Mai 1961 | Tiliaceae |
|  | *Intratriporopollenites cecilensis* Krutzsch 1961 | Tiliaceae |
| *Subtriporopollenites* spp. | *Subtriporopollenites anulatus* Thomson & Pflug 1953 *nanus* Thomson & Pflug 1953 | Juglandaceae |
|  | *Subtriporopollenites constans* Pflug 1953 | ? |
|  | *Subtriporopollenites supracirculus* Krutzsch & Vanhoorne 1977 | Juglandaceae |
| other triporate pollen | *Olaxipollis matthesi* Krutzsch 1962 | Olacaceae? |
|  | *Caryapollenites triangulus* (Pflug 1953) Krutzsch 1961 | Juglandaceae, C*arya* |
|  | *Celtipollenites intrastructurus* (Krutzsch & Vanhoorne 1977) Thiele-Pfeiffer 1980 | Ulmaceae, *Celtis* |
|  | *Celtipollenites laevigatus* Thiele-Pfeiffer 1988 | Ulmaceae |
|  | *Triporopollenites* spp*.,* 2 unidentified morpho-types | ? |
| *Alnipollenites verus* | *Alnipollenites verus* R. Potonié 1931 emend. R. Potonié 1960 | Betulaceae, *Alnus* |
| *Tricolpopollenites liblarensis* group | *Tricolpopollenites liblarensis* (Thoms. in R. Pot., Thoms. & Thierg. 1950) Th. & Pf. 1953 *liblarensis* (Thoms. in R. Pot., Thoms. & Thierg. 1950) Th. & Pf. 1953 | Fagaceae, Fabaceae, Combretaceae, Verbenaceae |
|  | *Tricolpopollenites liblarensis* (Thoms. in R. Pot., Thoms. & Thierg. 1950) Th. & Pf. 1953 *fallax* (R. Potonié 1934) Thomson & Pflug 1953 | Fagaceae, Fabaceae, Combretaceae, Verbenaceae |
|  | *Tricolpopollenites quisqualis* (R. Potonie 1934) Thomson & Pflug 1953 | Fagaceae? |
| *Tricolpopollenites retiformis* | *Tricolpopollenites retiformis* Thomson & Pflug 1953, 2 morpho-types | Salicaceae, *Salix* |
| *Tricolpopollenites vegetus* | *Tricolpopollenites vegetus* (R. Potonié 1934) Krutzsch 1959 | Hamamelidaceae? |
| *Tricolpo(ro)pollenites parmularius* | *Tricolpo(ro)pollenites parmularius* (R. Potonié 1934) Kr. in Kr., Pch. & Spieg. 1960 | Eucommiaceae, *Eucommia* |
| *Tricolporopollenites belgicus* | *Tricolporopollenites belgicus* Krutzsch & Vanhoorne 1977 | ? |
| *Spinaepollis spinosus* | *Spinaepollis spinosus* (R. Potonié 1931) Krutzsch 1961 | Euphorbiaceae? |
| *Tricolporopollenites cingulum* group | *Tricolporopollenites cingulum* (R. Potonie 1931) Thomson & Pflug 1953 *fusus* (R. Potonie 1931) Thomson & Pflug 1953, 2 morpho-types | Fagaceae, *Castanopsis*? |
|  | *Tricolporopollenites cingulum* (R. Potonie 1931) Thomson & Pflug 1953 *pusillus* (R. Potonie 1934) Thomson & Pflug 1953 | Fagaceae, *Castanopsis, Lithocarpus, Pasania* |
|  | *Tricolporopollenites cingulum* (R. Potonie 1931) Thomson & Pflug 1953 *oviformis* (R. Potonie 1931) Thomson & Pflug 1953 | Fagaceae, *Castanea*, *Castanopsis, Lithocarpus, Pasania* |
| *Nyssapollenites* spp. | *Nyssapollenites kruschii* (R. Pot. 1931) Nagy 1969 *analepticus* (R. Pot. 1934) Nagy 1969 | Nyssaceae, *Nyssa* |
|  | *Nyssapollenites kruschii* (R. Pot. 1931) R. Pot., Th. & Thierg. 1950 *accessorius* (R. Pot. 1934) R. Pot., Th. & Thierg. 1950 ex Simoncsics 1969 | Nyssaceae, *Nyssa* |
|  | *Nyssapollenites contortus* (Thomson & Pflug 1953) Nagy 1985 | Nyssaceae, *Nyssa* |
| *Ilexpollenites* spp. | *Ilexpollenites iliacus* (R. Pot. 1931) Thierg. 1937 ex 1937 ex R. Pot. 1960 | Aquifoliaceae, *Ilex* |
|  | *Ilexpollenites margaritatus* (R. Pot. 1931) Thierg. 1937 ex 1937 ex R. Pot. 1960 | Aquifoliaceae, *Ilex* |
| *Zonocostites* *ramonae* group | *Zonocostites ramonae* Germeraad, Hopping & Muller 1968 | Rhizophoraceae, *Rhizophora* |
|  | *Tricolporopollenites mansfeldensis* Krutzsch 1969 | Rhizophoraceae? |
|  | *Tricolporopollenites pseudomansfeldensis* Krutzsch 1969 |  |
| *Tetracolporopollenites* spp. | *Tetracolporopollenites sapotoides* Thomson & Pflug 1953 | Sapotaceae |
|  | *Tetracolporopollenites manifestus* (R. Potonié 1931) Thomson & Pflug 1953 | Sapotaceae |
| other tricolp(or)ate pollen | *Tricolp(or)opollenites staresedloensis* Krutzsch & Pacltová 1969 | Salicaceae? |
|  | *Tricolpopollenites constrictus* (Pierce 1961) Krutzsch & Vanhoorne 1977 | ? |
|  | *Tricolpopollenites asper* Thomson & Pflug 1953 | Fagaceae, *Quercus?* |
|  | *Tricolpopollenites* spp. – 7 unidentified morpho-types | ? |
|  | *Phimopollenites striolata* Hu, Jarzen & Dilcher 2008 | ? |
|  | *Tricolporopollenites euryoides* Kohlmann-Adamska and Ziembinska-Tworzydlo 2014 | Pentaphylacacaeae**,** *Eurya* |
|  | *Tricolporopollenites cf. baculatus* Krutzsch 1961 | ? |
|  | *Tricolporopollenites reticingulum* Krutzsch & Vanhoorne 1977 | ? |
|  | *Tricolporopollenites crassiexinus* Krutzsch & Vanhoorne 1977 | Celastraceae |
|  | *Tricolporopollenites exactus* (R. Potonie 1931) Thomson & Pflug 1953 | Cyrillaceae |
|  | *Tricolporopollenites megaexactus* (R. Potonie 1931) Thomson & Pflug 1953 *exactus* (R. Potonie 1931) Thomson & Pflug 1953 | Cyrillaceae |
|  | *Tricolporopollenites megaexactus* (R. Potonie 1931) Thomson & Pflug 1953 *brühlensis* (Th. in R. Pot., Th., & Thier. 1959) Thomson & Pflug 1953 | Cyrillaceae |
|  | *Tricolporopollenites marcodurensis* Thomson & Pflug 1953 | Vitaceae |
|  | *Tricolporopollenites megaporatus* Krutzsch & Vanhoorne 1977, 2 morpho-types | ? |
|  | *Tricolporopollenites solé de portai* Kedves 1965 | Anacardiaceae, Rosaceae |
|  | *Tricolporopollenites stiatopunctatus* Krutzsch & Vanhoorne 1977, 2 morpho-types | ? |
|  | *Tricolporopollenites abbreviatus* (R. Pot. 1934) Krutzsch 1961 | ? |
|  | *Tricolporopollenites eofagoides* Krutzsch & Vanhoorne 1977 | ? |
|  | *Tricolporopollenites magniferoides* Slodkowska 2014 | Anacardiaceae |
|  | *Tricolporopollenites satzveyensis* Pflug 1953 | Mastixiaceae |
|  | *Tricolporopollenites edmundii* (R. Potonié 1931) Thomson & Pflug 1953 | Mastixiaceae |
|  | *Tricolporopollenites pseudointergranulatus* Krutzsch 1966 ex Sontag 1966, 2 morpho-types | Menispermaceae |
|  | *Tricolporopollenites microreticulatus* Thomson & Pflug 1953 | Oleaceae |
|  | *Tricolporopollenites messelensis* Thiele-Pfeiffer 1988 | ? |
|  | *Tricolporopollenites* spp. – 6 unidentified morpho-types | ? |
|  | *Edmundipollis mastixioides* Slodkowska & Ziembinska-Tworzydlo 2014 | Mastixiaceae |
|  | *Verrutricolporites regillus* (R. Potonie 1934) Kedves 1978 | ? |
|  | *Parthenopollenites formosus* (Mamczar 1960) Worobiec 2014 | Vitaceae |
|  | *Spinulaepollis arceuthobioides* Krutzsch 1962 | ? |
|  | *Araliaceoipollenites euphorii* (R. Potonié 1931) R. Potonié 1951 | Araliaceae |
| *Ericipites* spp. | *Ericipites callidus* (R. Potonié 1931) Krutzsch 1970 | Ericaceae |
|  | *Ericipites ericius* (R. Potonié 1931) R. Potonié 1960 | Ericaceae |
| Taxa that have been used for diversity analysis, but which were, due to their low abundance, not used for pollen diagrams and multivariate statistics | *Polyporopollenites eoulmoides* Krutzsch & Vanhoorne 1977 | Ulmaceae, *Zelkova*? |
|  | *Polyvestibulopollenites juglandaceoides* Krutzsch & Vanhoorne 1977 | Juglandaceae |
|  | *Pterocaryapollenites stellatus* (R. Potonié 1931) Thiergart 1937 | Juglandaceae, *Pterocarya* |
|  | *Porocolpopollenites vestibulum* (R. Potonié 1931) Thomson & Pflug 1953 | Symplocaceae, *Symplocos* |
|  | *Porocolpopollenites rarobaculatus* Thiele-Pfeiffer 1980 | Symplocaceae, *Symplocos* |
|  | *Symplocospollenites orbis* (Thomson & Pflug 1953) R. Potonié 1960 | Symplocaceae, *Symplocos* |
|  | *Parsonsidites britannicus* Gruas-Cavagnetto 1976 | Malvaceae, Balanophoraceae?, Apocynaceae |
|  | *Compositoipollenites minimus* Krutzsch & Vanhoorne 1977 | ? |
|  | *Malvacipollis subtilis* Stover 1973 | Malvaceae?, Euphorbiaceae |
|  | *Erdtmanipollis pachysandroides* Krutzsch 1966 | Buxaceae, *Pachysandra* |
|  | *Reductipollis reductus* Krutzsch 1966 | Buxaceae? |
|  | *Polycolpites helmstedtensis* Krutzsch 1969 | Rubiaceae |
| **Algae** | | |
| *Apectodinium* spp. | *Apectodinium longispinosum* (Wilson 1968) Bujak & Davies 1983 | Dinoflagellata |
|  | *Apectodinium parvum* (Alberti 1961) Lentin & Williams 1977 | Dinoflagellata |
|  | *Apectodinium homomorphum* (Deflandre & Cookson, 1955) Lentin & Williams, 1977 | Dinoflagellata |
|  | *Apectodinium quinquelatum* (Williams & Downie 1966) Costa and Downie, 1979 | Dinoflagellata |
| *Botryococcus* | *Botryococcus* cf*. braunii* Kützing 1849 | Chlorophyceae |
| Zygnemataceae | *Gelasinicysta* sp*. (Zygnema-*type*)* | Zygnemataceae |
|  | *Ovoidites sp.* | Zygnemataceae |
|  | *Debarya-*type | Zygnemataceae |
|  | Unidentified zygospore | Zygnemataceae |
| other freshwater algae | *Planctonites stellarius* (R. Potonié 1934) Krutzsch 1960 | Chlorophyceae |
|  | *Tetraporina spinifera* Lindgren 1980 | Chlorophyceae |
|  | *Tetrapidites sp.* | Chlorophyceae |
|  | Unidentified zygospore | Chlorophyceae |
