## Supplementary material for "Greenhouse conditions in lower Eocene coastal wetlands? – Lessons from Schöningen, Northern Germany": S6_Table

**S5 Table: Estimations of beta diversity for Seam 1 in section S1.**

|  | S1-1 | S1-2 | S1-3 | S1-4 | S1-5 | S1-6 | S1-7 | S1-8 | S1-9 | S1-10 | S1-11 |
| --- | --- | --- | --- | --- | --- | --- | --- | --- | --- | --- | --- |
| S1-11 | 0.47 | 0.48 | 0.48 | 0.50 | 0.73 | 0.54 | 0.52 | 0.44 | 0.44 | 0.51 | - |
| S1-10 | 0.58 | 0.54 | 0.53 | 0.53 | 0.60 | 0.61 | 0.52 | 0.54 | 0.49 | - | 0.51 |
| S1-9 | 0.55 | 0.52 | 0.56 | 0.60 | 0.68 | 0.56 | 0.53 | 0.45 | - | 0.49 | 0.44 |
| S1-8 | 0.48 | 0.53 | 0.52 | 0.50 | 0.65 | 0.51 | 0.51 | - | 0.45 | 0.54 | 0.44 |
| S1-7 | 0.51 | 0.54 | 0.46 | 0.42 | 0.64 | 0.44 | - | 0.51 | 0.53 | 0.52 | 0.52 |
| S1-6 | 0.63 | 0.55 | 0.56 | 0.46 | 0.61 | - | 0.44 | 0.51 | 0.56 | 0.61 | 0.54 |
| S1-5 | 0.79 | 0.61 | 0.66 | 0.66 | - | 0.61 | 0.64 | 0.65 | 0.68 | 0.60 | 0.73 |
| S1-4 | 0.53 | 0.47 | 0.46 | - | 0.66 | 0.46 | 0.42 | 0.50 | 0.60 | 0.53 | 0.50 |
| S1-3 | 0.40 | 0.42 | - | 0.46 | 0.66 | 0.56 | 0.46 | 0.52 | 0.56 | 0.53 | 0.48 |
| S1-2 | 0.46 | - | 0.42 | 0.47 | 0.61 | 0.55 | 0.54 | 0.53 | 0.52 | 0.54 | 0.48 |
| S1-1 | - | 0.46 | 0.40 | 0.53 | 0.79 | 0.63 | 0.51 | 0.48 | 0.55 | 0.58 | 0.47 |

Given are pairwise comparisons of 11 lignite samples from section S1 using the measure of Whittaker (1960, 1972): (S/ā) – 1; S, total number of species in the two compared samples, ā, average number of species in the two compared samples of Seam 1.
